# The potential of plasma HLA peptides beyond neoepitopes

**DOI:** 10.1101/2023.09.05.556309

**Authors:** Maria Wahle, Marvin Thielert, Maximilian Zwiebel, Patricia Skowronek, Wen-Feng Zeng, Matthias Mann

**Affiliations:** Department Proteomics and Signal Transduction, Max Planck Institute of Biochemistry, Martinsried, Germany

## Abstract

Distinction of non-self from self is the major task of the immune system. Immunopeptidomics studies the peptide repertoire presented by the human leukocyte antigen (HLA) protein, usually on tissues. However, HLA peptides are also bound to plasma soluble HLA (sHLA), but little is known about their origin and potential for biomarker discovery in this readily available biofluid. Currently, immunopeptidomics is hampered by complex workflows and limited sensitivity, generally requiring several mL of plasma for the detection of hundreds of HLA peptides. Here, we take advantage of recent improvements in the throughput and sensitivity of mass spectrometry (MS)-based proteomics to develop a highly-sensitive, automated and economical workflow for HLA peptide analysis, termed Immunopeptidomics by Biotinylated Antibodies and Streptavidin (IMBAS). IMBAS-MS quantifies more than 5,000 HLA class I peptides from only 200 μL of plasma, in just 30 minutes. Our technology revealed that the plasma immunopeptidome of healthy donors is remarkably stable throughout a year and strongly correlated between individuals with overlapping HLA types. Immunopeptides originating from diverse tissues, including the brain, are proportionately represented. We conclude that sHLAs are a promising avenue for immunology and precision oncology.

## INTRODUCTION

The immune system relies on the human leukocyte antigen (HLA) peptide-protein complex to present immunopeptides derived from both endogenous and exogenous sources to circulating T-cells. These immunopeptides play a crucial role in immune surveillance, as they enable elimination of abnormal or infected cells. Upon recognition of a peptide-HLA protein complex by cytotoxic T lymphocytes (CTLs), downstream cascades are activated, causing the presenting cell to undergo apoptosis. This biological principle is exploited in immunotherapeutic strategies such as CAR T-cell treatment or mRNA peptide vaccination^1^. A crucial but challenging step is the identification of peptides specifically presented by tumor cells. Most efforts have focused on the enrichment of membrane bound human leukocyte antigen (mHLA) receptors with their bound immunopeptides from tumor tissue. This is followed by mass spectrometric identification in search of tumor specific antigens or neoepitopes^2^,^3^.

In addition to the membrane anchored HLA proteins, a soluble fraction (sHLA) can be enriched from blood and other body fluids^4^. sHLAs are thought to arise by shedding, cleavage from the cell surface or the expression of a splicing variant lacking the membrane anchoring domain^5^. Although their function and exact release mechanisms remain unclear, although it is known to change in a disease context^6^. In the case of cancer, a disproportionate fraction of peptides in relation to the total tumor mass may originate from tumor tissue^4^, representing a potential additional source for tumor antigens^7^. Whereas the analysis of HLA peptides from mHLAs requires substantial tissue amounts from surgery, sHLAs are rather easily accessible through minimal invasive procedures like a regular blood withdrawal without placing additional burden on patients. However, despite the clear potential for disease diagnosis and treatment monitoring in a clinical setup^8–10^, there are only few studies investigating sHLAs and they typically only identified hundreds of them from milliliters of plasma^4,7,8^.

Beyond clinical applications, another attraction of the sHLA immunopeptidome is that it can serve as an unlimited source of native immunopeptides from diverse HLA backgrounds. As importantly, an extensive repertoire of sHLA peptides from a diversity of healthy donors could potentially serve as a resource to improve general knowledge about peptide processing and presentability^11^. In contrast, due to limited analytical sensitivity and tissue accessibility mHLA data is currently restricted to a few hundred different alleles, with on average around 5,000 up to 160,000 (HLA-A0201) associated MHC peptides identified by mass spectrometry^12^.

In this study, we address the limitations of sHLA immunopeptidomics and characterize its nature over time in healthy donors. We describe an automated workflow for efficient ‘one-pot’ enrichment of HLA immunopeptides followed by ultra-high sensitive mass spectrometry (MS), termed IMBAS-MS for Immunopeptidomics by Biotinylated Antibodies and Streptavidin. Quantifying immunopeptides from a few hundred microliters showed that the immunopeptidome is stable in healthy individuals for over a year and exhibits high reproducibility between overlapping HLA types. IMBAS-MS allowed profiling of sHLA to a depth of over 10,000 peptides per person, which were broadly representative of different tissues.

## EXPERIMENTAL PROCEDURES

### Plasma collection from healthy donors

Plasma was obtained by withdrawing blood from eight healthy donors (specified in Supplementary Table 1) in EDTA tubes (BD Vacutainer K2E; REF 367525). The tubes were inverted three times and centrifuged at 2000xg for 20 min at 4°C. The plasma was separated, aliquoted and snap-frozen and stored at -80°C.

To determine the HLA types of healthy donors, genomic DNA was extracted from buccal swabs. Cotton swabs were used to obtain oral mucosa samples, placed in protease buffer (30 mM Tris-HCl, pH 8, 0.5% Tween-20, 0.5% IGEPAL CA-630) and proteins were digested with 100 μg proteinase K per sample for 12 min at 50°C in a thermal shaker. The enzyme was inactivated at 75°C for 30 min and genomic DNA was purified using the QIAamp DNA Micro Kit (Qiagen). HLA class I (HLA-A, HLA-B, HLA-C) and class II (DRB1, DQB1, DPB1) loci were amplified using the NGSgo®-MX6-1 kit (GenDx) and multiplexed sequencing libraries were prepared using the Ultra™ II FS DNA Library Prep Kit for Illumina (NEB). Libraries were sequenced on a NovaSeq 6000 system (Illumina) in 150 bp paired-end mode and the genotype data were analyzed using the NGSengine® software (GenDx). Blood was sampled from healthy donors, who provided written informed consent, with prior approval of the ethics committee of the Max Planck Society.

### Affinity purification of HLA Molecules

Plasma samples were thawed on ice, diluted 1:2 with PBS (Gibco) and incubated overnight, shaking at 4°C with variable amounts of biotinylated W6/32 antibody (custom produced by inVivo Bioscience). Captured HLA molecules were enriched using magnetic streptavidin beads (ReSyn Bioscience) and washed first with 100 μl of 150 mM HCl in 10mM Tris pH8.5, then 100 μl of 450 mM HCl in 10 mM Tris pH 8.5 and finally 100 μl of 10 mM Tris pH 8.5 at 4°C. The sHLA molecules were eluted from the beads using 150 μl elution buffer (200 mM glycine pH 2), transferred into prewetted 30 kDa MWCO plates (Millipore) and filtered at 4000xg for 20 min. The flowthrough was directly loaded onto Evotips Pure following the recommended standard procedure. Briefly, Evotips were activated by 1-propanol, washed two times with 50 μL buffer B (99.9% ACN, 0.1% FA) and two times with 50 μL buffer A (99.9% ddH2O, 0.1% FA). 70 μL buffer A was briefly spined on the disks and sample elution was loaded by 80 sec centrifugation. Evotips were then washed with 50 μL buffer A and stored with buffer A on top. All centrifugation steps were performed at 700 x g for 1 min. The whole protocol was performed in a semi-automated fashion using the Agilent Bravo liquid handling platform.

### Immunopeptide fractionation

To acquire deep fractionated immunopeptidomes, 5 ml of plasma per individual were thawed at once, distributed in 10 wells of a 96 deep-well plate and processed with the same workflow as described above. The final elutions were pooled in two wells before loading onto the MWCO filter plate and combined into one single peptide pool after centrifugation. Fractionation was carried out on an AssayMAP Bravo Sample Prep Platform (Agilent), using the Fractionation v1.1 Protocol in the Protein Sample Prep Workbench v3.2.0 with standard settings. 5 μl C18 Cartridges (Agilent) were used as solid phase, six elution fractions were collected, using high-pH buffers with increasing acetonitrile concentrations (ammonium hydroxide solution, pH 10; 7, 12, 15, 23, 30 and 40% acetonitrile, respectively). The 40% acetonitrile buffer was used for priming and 7% for equilibration of the cartridges.

### DDA and DIA LC-MS acquisition

Peptides were separated with the Evosep One LC system using predefined gradients as mentioned in each section. The majority of data was acquired using the Whisper40 method over an 15 cm Aurora Elite CSI column (AUR3-15075C18-CSI, IonOpticks) at 50°C C inside a nanoelectrospray ion source (Captive spray source, Bruker). The mobile phases were 0.1% FA in LC-MS grade water (buffer A) and 99.9% ACN with 0.1% FA (buffer B). For gradient testing, the 15SPD, 30SPD and 60SPD method was used in combination with a 15 cm PepSep (150 um ID and 1.5 um bead size, Bruker) connected to a 10um ID ZDV emitter (Bruker). The LC system was coupled to a timsTOF Ultra instrument (Bruker).

When operated in dda-PASEF mode, a ten PASEF/MSMS scan per topN acquisition method was used with a precursor signals intensity threshold at 500 arbitrary units. An adapted polygon in the m/z-IM plane was used to exclude adverse ions, but include single-charge precursors. The mass spectrometer was operated in sensitivity mode with an accumulation and ramp time of 100ms. Precursors were isolated with a 2 Th window below m/z 700 and 3 Th above and actively excluded for 0.4 min when reaching a target intensity threshold of 20,000 arbitrary units. A range from 100 to 1700 m/z and 0.6 to 1.6 Vs cm-2 was covered with a collision energy from 20 eV at 0.6 Vs cm-2 ramped linearly to 59 eV at 1.6 Vs cm-2.

When operating in dia-PASEF mode, we used optimal dia-PASEF methods generated with our Python tool py_diAID^30^. These dia-PASEF methods optimally cover the precursor cloud in m/z-IM plane, while being highly efficient with 1.17 s cycle time. We generated acquisition schemes specifically for dominant HLA types that cover up to 99.9% of all precursor species including singly charged ions, with 8 dia-PASEF scans, where each scan is divided into two ion mobility windows. The method covers precursors within 300 – 1200 Da. Other settings remained the same as for dda-PASEF.

### Raw data analysis

DDA data was analyzed using FragPipe 19.1 with the nonspecific HLA workflow^19–21^. The quality of identified peptides was assessed using MHCVizPipe (v0.7.11) ^31^.

DIA data was analyzed using DIA-NN version 1.8.1 with standard settings searching against sample specific predicted libraries generated using the AlphaPeptDeep package (https://github.com/MannLabs/alphapeptdeep) together with the peptdeep_hla (https://github.com/MannLabs/PeptDeep-HLA) DL model. Sample specific immunopeptide lists (from DDA analysis or directDIA results generated using Spectronaut 17) were used to tune an immunopeptide deep learning model which reports a peptide list with high likelihood of being a presented immunopeptide in this sample from an unspecific *in silico* digest of a human fasta. The peptide list of each individual is used for transfer learning to predict individual- or sample-specific spectral library. Library sizes are adjustable using a precision cutoff (probability >= 0.7) in the peptdeep_hla DL model. The ‘report.tsv’ table was used for further analysis.

### Statistical analysis

All data analysis was performed using R. Unless stated differently, only peptides predicted to bind to any of a donors HLA-type were used for downstream analysis. Peptides were defined to be binders as provided by MHCVizPipe interfacing NetMHCpan 4.1. Binder Frequency (BF) scores were reported as provided by MHCVizPipe. The BF score describes the fraction of peptides predicted to bind the provided HLA Alleles within the expected length range. The UpSet plots were generated using a custom script. Only intersections which have a size of 20% of the smallest included dataset are displayed.

### Experimental design and statistical rational

All experiments were done using human plasma obtained as described above. Altogether, the dataset including raw data files and search results were uploaded to MassIVE (see below). We used the same plasma batch for the benchmarking and technical evaluation. In brief, measurements with different gradients, input amounts or from different donors were acquired in triplicates unless mentioned differently. The experimental design and statistical rational are described in the respective figure legends. Workflow replicates were acquired to evaluate reproducibility and quantitative accuracy.

## RESULTS

### IMBAS-MS design and evaluation

Earlier sHLA workflows have used several milliliters of plasma for enrichment, limiting throughput and applicability. We aimed to develop a workflow that is highly reproducible, sensitive and allows for deep immunopeptidome coverage, without neglecting throughput and cost. To accommodate all these aspects, we optimized IMBAS (Immunopeptidomics by Biotinylated Antibodies and Streptavidin) in a 96-well format that could be processed in parallel. We automated the immunoaffinity enrichment on a Bravo liquid handling robot (Agilent) with less than two hours of hands-on time for the entire procedure (Fig. 1A). The workflow was designed to be flexible, thus the enrichment can either be performed by hand or by any robot with a magnetic plate and a cooled plate station. IMBAS-MS is modular and although not demonstrated here, can directly be applied to cell lysates or biopsy samples by adding a homogenization and lysis step up front. The enrichment, washing and elution steps take place within the same well, minimizing transfer steps and reducing sample loss due to plastic contact. A key aspect of IMBAS is the replacement of the standard ProteinA/G-IgG domain interaction between the antibody and bead matrix. To achieve this, we chose to use biotinylated antibodies which can be captured with streptavidin beads. The high specificity and stability of the streptavidin-biotin interaction allows to omit chemical crosslinking of the desired anti-HLA-antibody to the slurry upfront of the enrichment protocol, saving time and material. Additionally, this eliminates the plasma preclearance step. Following the enrichment, the eluent is molecular-weight filtered and the resulting, separated peptides are loaded onto Evotips. In this way, an entire 96-well plate can be prepared and MS-data acquired within three days with minimal reagent preparation and cost. For technical details of the protocol see Experimental Methods.

**Figure 1:**
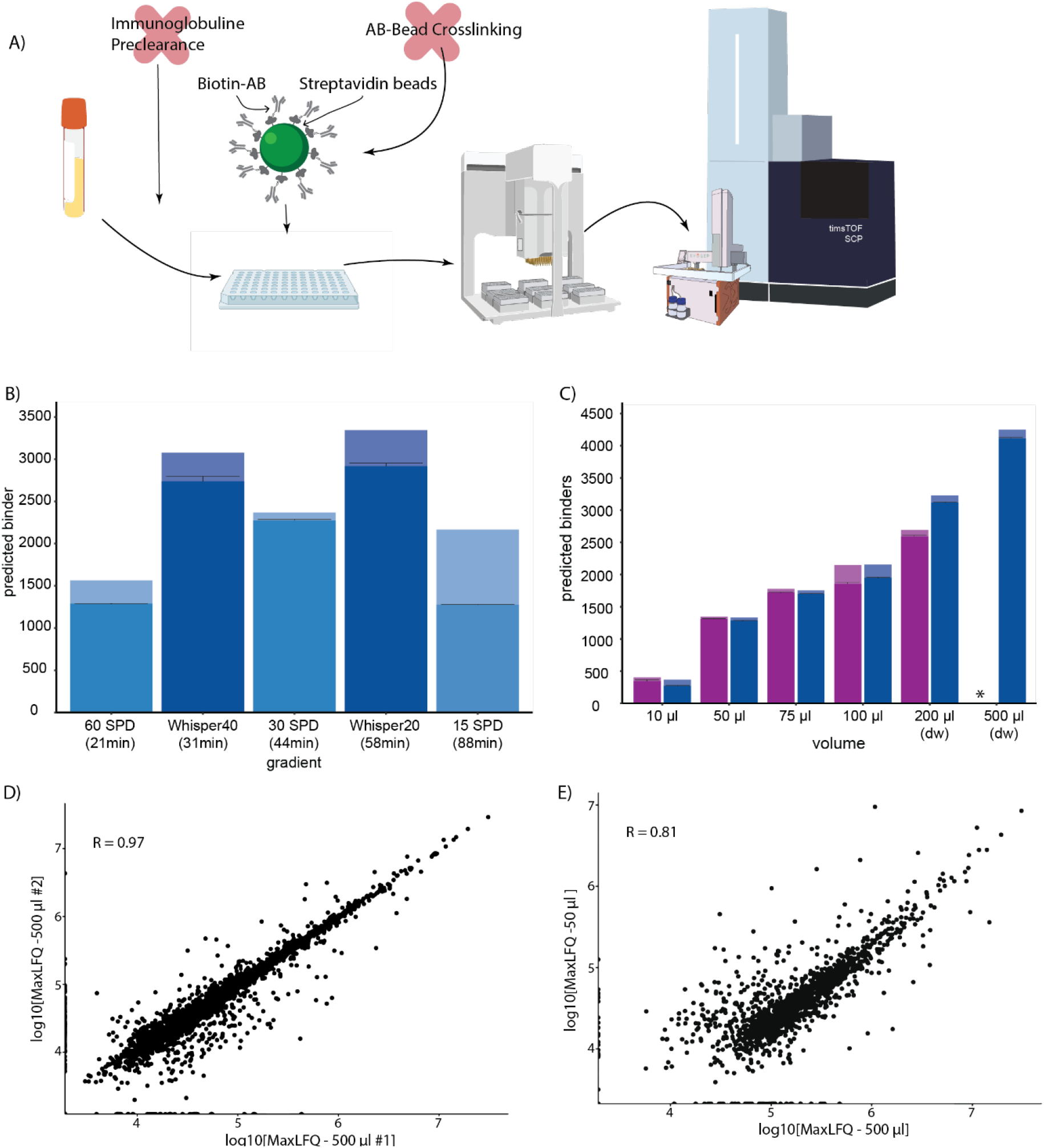
IMBAS-MS design and evaluation. **(A)** Schematic representation of the sHLA IMBAS-MS workflow. Plasma is incubated with anti-HLA antibody (W6/32) and subjected to our automated bead-based enrichment workflow on a liquid handling robot (Agilent Bravo). Eluted peptides are loaded onto StageTips (Evotips) and measured by ultra-high sensitivity LC-MS/MS (Evosep and Bruker timsTOF Ultra). **(B)** Immunopeptide identifications from enrichment of 200 μL plasma using different gradient types and lengths. Low-flow gradients (Whisper40 and 20 on the Evosep, dark blue) show an increased sensitivity by identifying over 3,000 immunopeptides. The total uniquely identified immunopeptides per triplicate (light bar) as well as the average and standard deviation (SD) per replicate (black lines) is plotted. **(C)** Evaluation of various input amounts using 1 ug (purple) or 10 ug (blue) of W6/32 antibody. The ‘*’ indicates that this data point was not acquired due to plasma/antibody requirements. For unique peptide number determination see B); dw: deep-well plate. **(D)** Quantification correlation shows high reproducibility between workflow replicates using 500 μL plasma. **(E)** Pearson correlation between 50 μL and 500 μL of the same plasma sample.

To evaluate IMBAS-MS, we first identified and quantified immunopeptides from 200 μL plasma from the same donor at different HPLC flowrates and gradient lengths, from 21 min up to 88 min long gradients (termed 60 to 15 Samples Per Day (SPD)). The standard methods on the Evosep system have 1 μL/min down to 0.22 μL/min flowrates with throughputs of 15, 30 and 60 SPD. We reasoned that the recently introduced very low flow gradients of only 100 nL/min (Whisper gradients20 or 40) that had substantially boosted sensitivity for single-cell analysis^13^ would also be beneficial for HLA peptides. Indeed, the nanoflow gradients substantially outperformed the standard gradients with more than 3,000 immunopeptide identifications in data-dependent acquisition (DDA) (Fig 1B). The Whisper20 gradient identified only 10% more peptides than the Whisper20 gradient, at the cost of doubled measurement time (Fig. 1B). With a focus on maximizing depth and throughput, we chose the Whisper40 gradient (31 min length) for all subsequent experiments.

A key advantage of our workflow is that it needs much less plasma input than the milliliters used before. To investigate input requirements and tradeoffs, we enriched sHLA from only 10 μL up to 500 μL of plasma. Volumes from 10 μL to 100 μL plasma can be processed in a standard 96 well plate. They required only 1 μg of antibody for efficient enrichment and did not benefit from increasing the antibody amount 10-fold (Fig. 1C). For higher volumes – for instance 200 μl – higher antibody amounts boosted immunopeptide identifications about 20%. Over the entire tested input range, we identified from 500 to 4,500 immunopeptides in data dependent acquisition (DDA).

To investigate the purity of our immunopeptidomes, we inspected their length distribution, their calculated binder scores and presence of singly charged precursors. Identified peptides retained expected features such as a strong preference of nonameric peptides and a significant proportion of singly charged precursors (Supplementary Figure 1). The fraction of peptides with high binding scores (BF) within the expected length range was 0.9, further indicating high purity of the enriched and identified peptides (see Experimental Methods).

IMBAS-MS also demonstrates high quantitative reproducibility between replicates at 500 μL (Pearson correlation of 0.97). Even with ten-fold reduced input, reproducibility is largely retained with a Pearson correlation of 0.81 (Figure 1D and E).

Based on the results above, in particular the purity and depth of the immunopeptide fraction, we chose a sample volume of 200 μL as an optimal combination for data quality, sample availability, ease of handling and cost effectiveness.

### Predicted library-based DIA for immunopeptidomics

Having evaluated IMBAS-MS with data dependent acquisition (DDA) methods, we set out to couple it with data-independent acquisition (DIA) based mass spectrometry, which promises much greater depth and higher data completeness between experiments^14,15^. A major challenge for efficient and comprehensive analysis of DIA data is the selection or generation of a suitable spectral library. Three different strategies are commonly used: experimental libraries, typically acquired by DDA; pseudospectra-based libraries extracted by directDIA as introduced by DIA-Umpire^16^ and implemented in Spectronaut; and libraries in which fragment intensities are predicted by deep learning^14,17^. In connection with the latter approach, we recently introduced a deep learning based framework called AlphaPeptDeep, which predicts spectral libraries tailored for different MS platforms, only based on a database file of the proteome in FASTA format or just a peptide list as input^18^. It contains the PeptDeep-HLA model which makes use of the inherent similarity of immunopeptides present within one person based on their HLA type. Given a preliminary list of identified peptides, this package then predicts a large subset of HLA peptides that are potentially present in this allelotype(s). Here, we compared three different modes of library generation in AlphaPeptDeep, purely experimental libraries and pseudospectra-extracted libraries (Fig. 2A).

**Figure 2:**
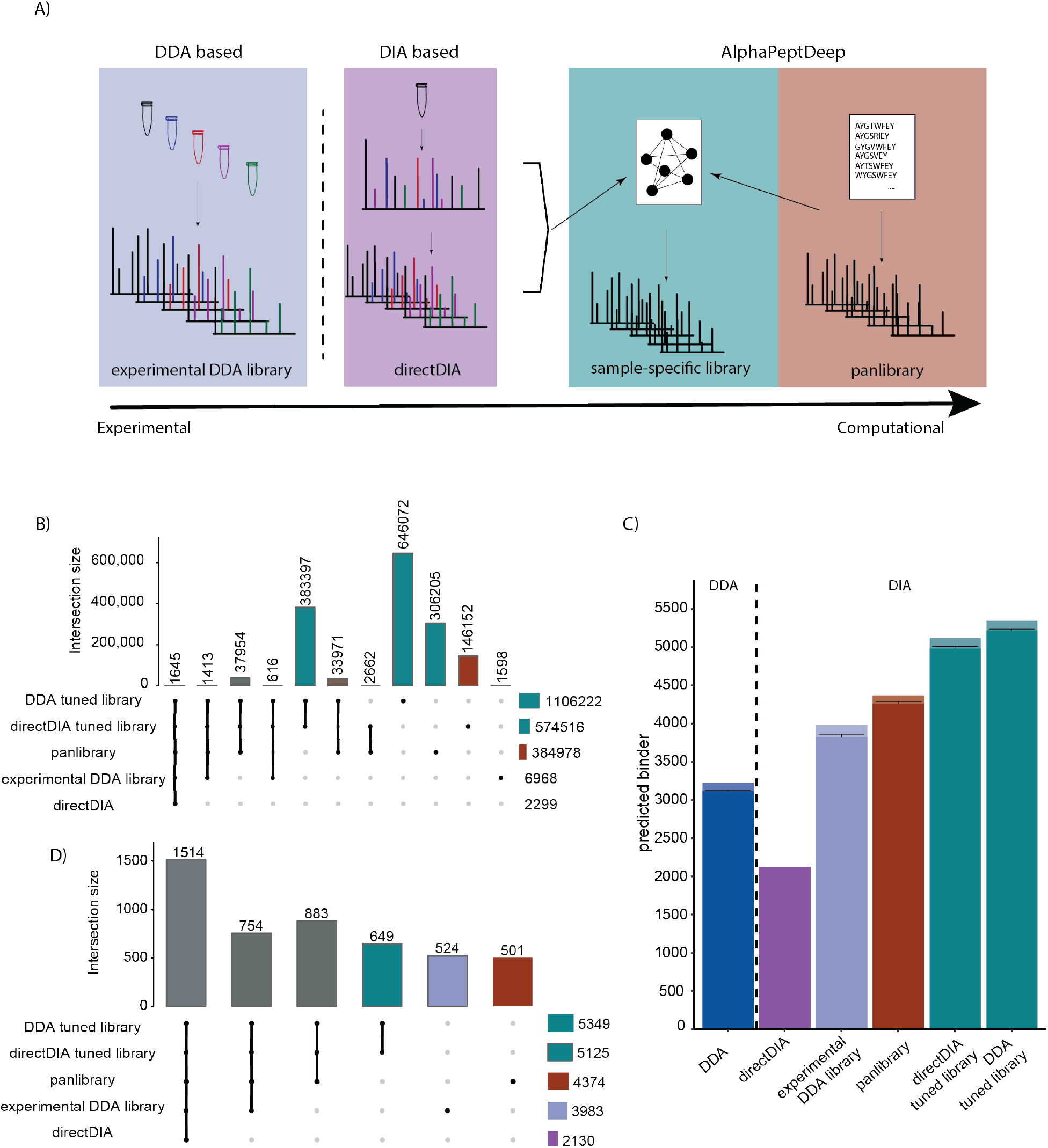
Evaluation of different strategies of predicted library-based DIA. **(A)** Overview of the library generation process. **(B)** Overlap of experimental library, pseudospectra library (directDIA), pan library and AlphaPeptDeep predicted libraries. **(C)** Triplicate measurement of immunopeptides from 200 μL plasma analyzed using different DIA data analysis strategies. DDA identifications are shown for comparison (left of the stippled vertical line). Library matching strategies are described in the main text. The height of the bar represents overall uniquely identified immunopeptides per triplicate and the mean per run as a horizontal black line with standard deviation. **(D)** Overlap of identified peptides analyzed with different library strategies (C).

First, we built an experimental DDA library using MSFragger^19–21^ on the above-described dilution series files, which resulted in nearly 7,000 identified precursors. Spectronaut internally builds a directDIA extracted list of peptides, in this case containing around 2,000 identified precursors from the replicates searched in parallel. Using AlphaPeptDeep, we predicted the fragment intensities of the set of immunopeptides constituting the ‘pan library’, with around 385,000 precursors (MSV000084172;PXD004894). The PeptDeep-HLA model only needs about 1,000 identified immunopeptides to learn how to extract potential immunopeptides from a FASTS for the allelotypes in question. For this, we compare two strategies, using either peptides identified in DDA experiments or from a directDIA search of the same file. On this basis, AlphaPeptDeep generated two large libraries (about 1 and 0.5 million precursors, respectively). Figure 2B compares the different DIA library sizes and their overlap.

With the exception of the directDIA strategy as currently implemented in Spectronaut, DIA always substantially outperformed DDA, as expected (Fig. 2C). Although a directDIA based analysis strategy as a standalone solution is not able to outperform a DDA immunopeptidomics analysis, a combination of directDIA with AlphaPeptDeep increased the depth by up to 67% compared to the DDA experiment (Fig 2C). Importantly, this increase in depth did not come at the expense of the quality of the data, as judged by the peptide length distribution and the binder scores which ranged from 0.9 for the experimental library to 0.98 for the DDA tuned library (Supplements). All three computational libraries, the pan library as well as the sample specific libraries outperform the experimental library while retaining the vast majority of peptides (Fig 2D). Given the small difference between the results from predicting the library based on DDA data or extracted peptides by directDIA, we suggest that the latter strategy will be attractive for immunopeptidomics in the future as no additional DDA experiments are required any more. We conclude that the combination of directDIA and AlphaPeptDeep enables us to acquire deep DIA based immunopeptidomes from a single measurement of a sample.

### Deep soluble immunopeptidomics in comparison to mHLA immunopeptidomics

Previous state of the art reports used around 10 mL plasma per donor to reach a median depth of around 1,000 unique immunopeptides with a maximum of 2,500 for healthy donors^7–9,22^. With our sample specific predicted spectral libraries and DIA acquisition IMBAS-MS, surpassed those results with just 2% of the input material (200 μL of plasma) (Fig. 3A). From one blood withdrawal that yielded 5 ml of plasma, we quantified up to 13,000 immunopeptides from six fractions per individual, six-fold higher than before (Fig 3A). (Note that the previous studies employed Q Exactive instruments rather than the latest generation Bruker timsTOFs.)

**Fig. 3:**
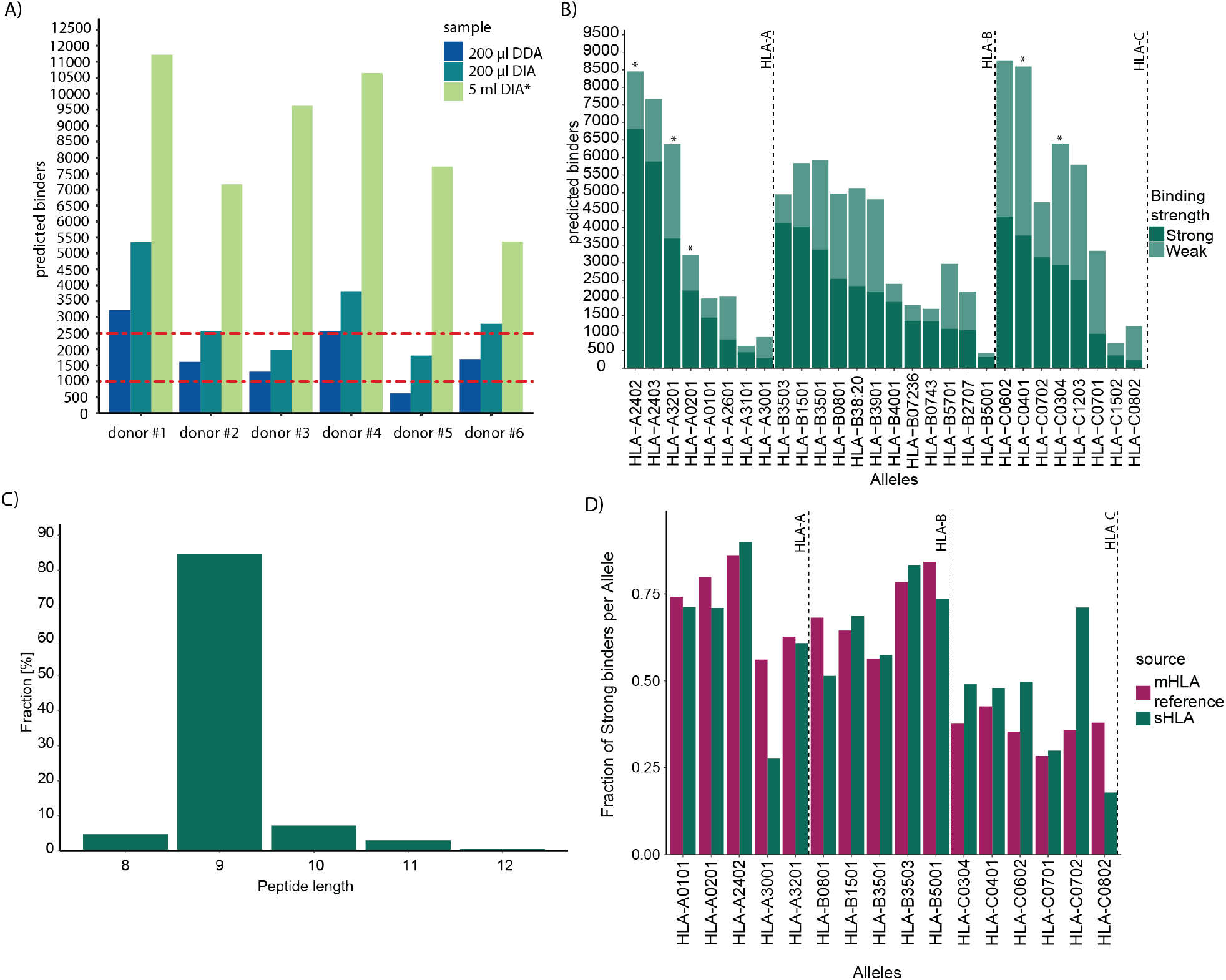
Deep soluble immunopeptidome retains properties described for mHLA based immunopeptidomes. **(A)** Immunopeptides identified by IMBAS-MS from 200 μL plasma by DDA (blue), DIA (green) or six separately analyzed fractions from 5 ml plasma (light green) from six healthy donors. For comparison the stippled red lines indicate the maximum and median identifications from 10 mL of ‘non-cancerous plasma’^7^.* The asterisk highlights that the 5 ml DIA runs were measured in six fractions. **(B)** Strong or weak predicted binders from (A) across HLA-types. Asterisks mark types present in multiple people. **(C)** Length distribution of identified HLA class I peptides of all samples in (A). **(D)** The fraction of strong binders for each allele present in our soluble HLA dataset from (A) compared to the fraction reported in a reference mHLA dataset (HLA-Ligandatlas^2^).

In total, the experiment above covered nearly 40,000 unique immunopeptides from 28 different alleles in six donors (Fig 3B). This is a large number, as evidenced by the fact that in some cases (e.g. HLA-C0401) this data surpasses the number of epitopes reported by the community database Immune Epitope Database & Tools (IEDB) which for this allelotype contains around 5,000 peptides identified by mass spectrometry compared to our 8,000 (https://www.iedb.org/result_v3.php?cookie_id=13cd62).

Next, we used our in-depth data to assess whether the soluble immunopeptidome displayed similar physicochemical properties as those described for their membrane bound equivalents. We noted that immunopeptides identified from sHLA show a strong enrichment of nonameric peptides, similar to what is known for immunopeptides from membrane bound equivalents (Fig 3C).

Comparing the ratio of peptides which bind strongly (%Rank <0.5) to their cognate HLA protein to those that bind weakly (0.5<%Rank<2), we did not find significant differences in pairwise comparisons of HLA types present in our sHLA dataset or an mHLA dataset^2^ (Fig. 3D). Given the easy accessibility of the sHLA peptidome by IMBAS-MS and its close correspondence to the mHLA peptidome, we conclude that plasma is a very attractive source to increase our general knowledge of immunopeptides.

### Tissue origin of sHLA peptides

A fundamental and still an open question is to what degree each organ contributes to the sHLA peptidome and we reasoned that our deep and unbiased dataset on healthy donors could shed light on this. To infer the origin of the soluble immunopeptidome we compare our immunopeptidome data to a recent and deep proteome atlas of 29 healthy tissues^23^. In that proteome dataset, each gene was classified into one of four groups, namely (i) expressed in all tissues, (ii) group enriched, (iii) tissue enhanced and (iv) tissue enriched. We transferred these classifications to our deep, fractionated sHLA peptidome to assign organ specificity to it. Next, we compared the frequency of group enriched, tissue enhanced and tissue enriched genes represented in the immunopeptidome to the frequency of those groups within the proteome of each tissue. Note that this assumes that proteins expressed in different organs have a similar chance to be presented by HLA proteins. That would make the fraction of genes assigned to each organ within the immunopeptidome a good estimate of the overall representation of that organ. We observed that the median frequency of classified genes in our immunopeptidome dataset correlates well with the frequency of those within the organ proteome dataset (R^2^ between 0.8 and 0.84). As an example, around 7.5% of all genes identified in the duodenum proteome were classified as group enriched^23^. This is very close to the value of 8% of all genes in our soluble immunopeptidomes. As can be seen in Figure 4A, ‘tissue enriched genes’ are less frequent in the proteome and in the immunopeptidome than genes belonging to the two less enriched groups. A notable exception from the above general observation are brain enriched genes. They are represented at around 3% within the immunopeptidomes, while encompassing around 5% of the brain proteome (Fig 4A). We speculate that sHLA-protein-peptide complexes may be partially filtered by the blood brain barrier or that brain specific genes are somewhat less likely to be presented, perhaps due to slow protein turnover. Another reason could be a lack of peptides originating from those proteins that have an affinity to one of the analyzed alleles in our dataset. In addition, we assessed whether the immunopeptidome quantified from the measurement of only 200 μL has sufficient depth to infer organ specific gene ontology enrichment terms. Indeed, applying the above gene classification strategy allows to discern organ function specific gene sets represented by immunopeptides as illustrated for donor#1 for liver enriched genes and brain enriched genes (Fig 4 B).

**Figure 4:**
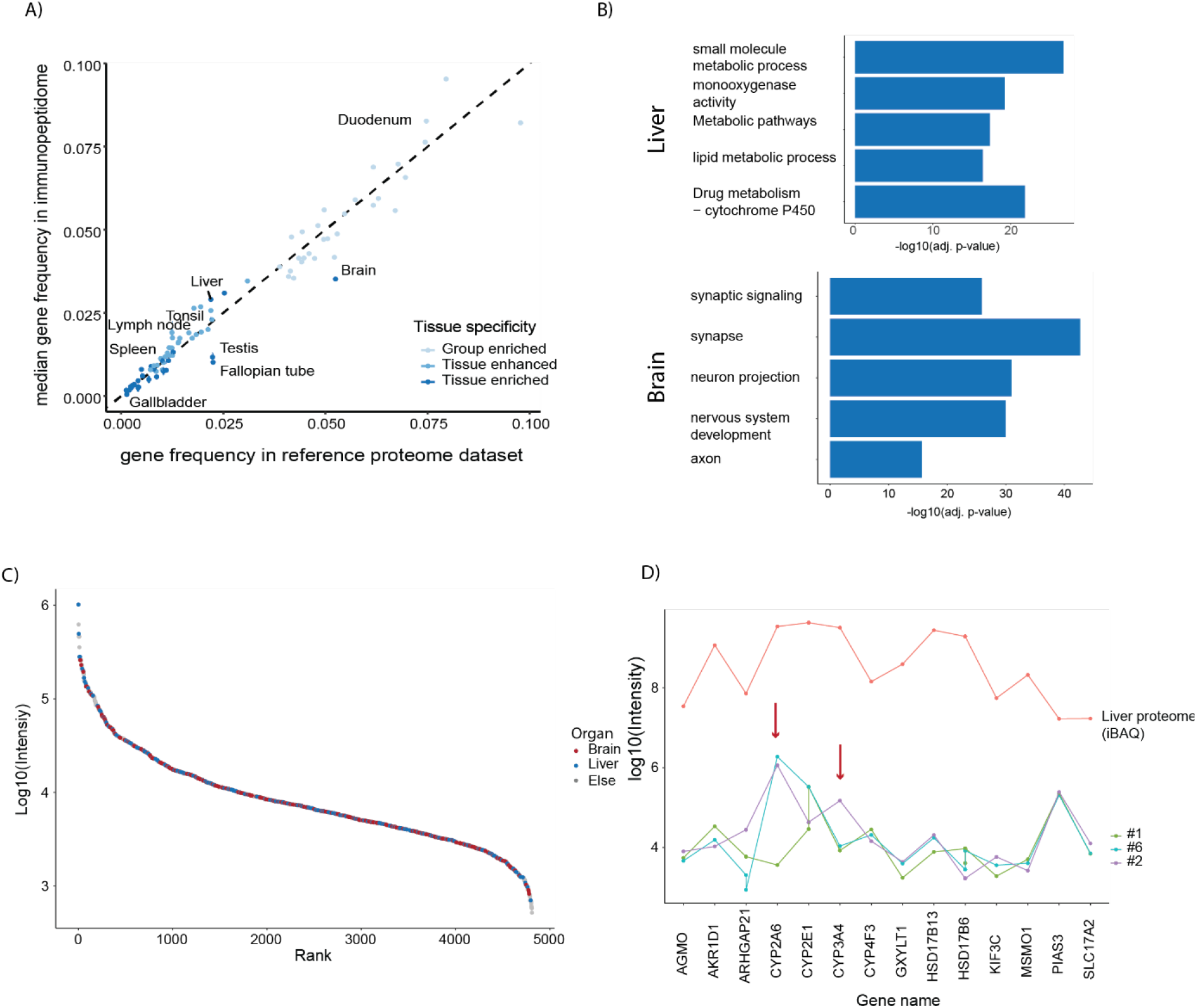
Tissue origin of immunopeptides. **(A)** Comparison of the fraction of immunopeptides assigned to different organs as inferred from a reference proteome dataset^23^. Genes and immunopeptides are classified as group enriched, tissue enhanced and tissue enriched depending on their degree of enrichment in the corresponding proteins reference proteome. The immunopeptidome dataset is the same as in Figure 3A. Points on or close to the diagonal indicate that the organ is equally represented in the peptidome and the proteome. **(B)** GO-term enrichment of immunopeptides whose proteins were assigned as tissue enhanced in brain or liver from donor #1 (dataset from Figure 3A, 200 μL DIA). Terms are representative of tissue specific functions. **(C)** Intensity rank plot of peptides derived from group enriched, tissue enhanced and tissue enriched genes colored by their respective organ assignment (dataset as in (B)). **(D)** Median intensity traces of selected liver tissue enriched immunopeptides for three donors. The iBAQ intensities for the corresponding genes from the reference proteome dataset are plotted for reference.

Interestingly in view of the different sizes of the organs, on a quantitative level, peptides representing all organs are distributed equally over the peptide abundance range with no clear trend of specific organs being represented by more high abundant species or vice versa. For example, both brain and liver assigned immunopeptides cover all three orders of magnitude in peptide intensities (Fig 4C, other organs see Supplementary Figure 2).

Among the set of tissue enriched liver genes common to three donors with shared HLA types, the trend of immunopeptide intensities is generally similar (Fig. 4D). However, there are clear differences in the presentation of Cytochrome-P-Oxygenase 2A6 (CYP) and CYP3A4, which are involved in the metabolization of nicotine and pharmaceutical drugs, respectively (red arrows). This may be attributable to lifestyle differences or genetic differences between donors.

Our results demonstrate that the soluble immunopeptidome is overall representative of the organs constituting the body. They also suggest a considerable potential of plasma immunopeptidome analysis for studying system-wide changes in the human proteome and in providing novel insights into physiological and pathological processes that are presented to the immune system.

### sHLA immunopeptidomic reproducability over time and between donors

While the immunopeptidome is thought to change considerably upon disease, little is known about its stability in healthy persons over time. To address this fundamental question, we followed an initially healthy person over a year. We sampled plasma at and shortly after the initial time point (16h apart) to gauge short term biological variation, at the five-month mark and at the end of the year. At about 11 months, the donor contracted COVID-19, and we sampled their immunopeptidome as soon as they were not positive any more (Supplementary Table 1).

Throughout the entire time period more than half of all sHLA peptides were detectable and quantifiable, with 88 to 93% being shared in at least two timepoints (Fig. 5A). Remarkably, quantitative reproducibility over the entire year was very high (Pearson correlation of 0.97 between the first and the last timepoint (Fig. 5B)). The first two sampling points that were only 16h apart, also agreed very well with each other, suggesting that time of day did not have a large influence. Even the immunopeptidome shortly after COVID-19 infection did not show large variations at a global scale.

**Figure 5:**
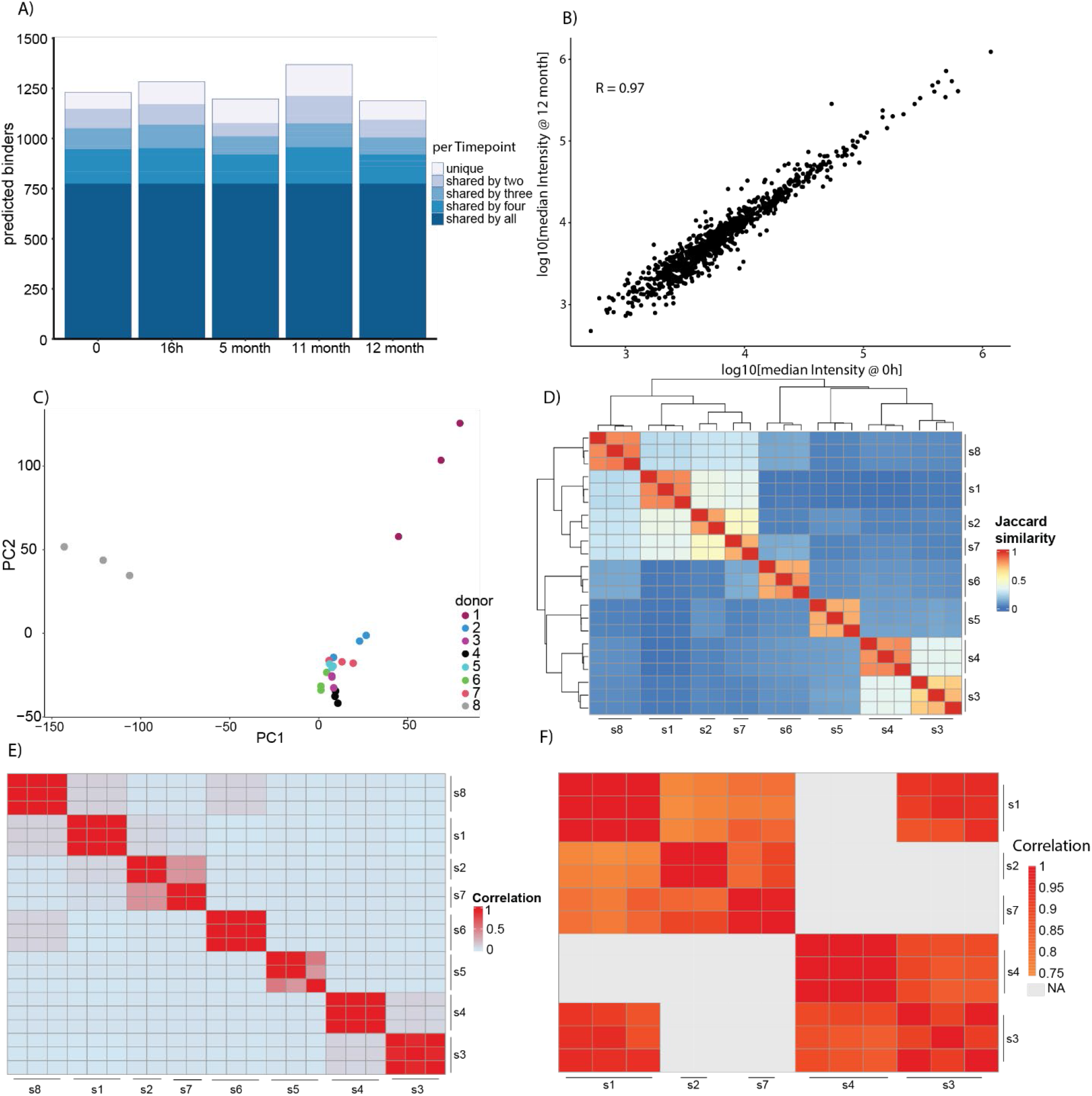
The plasma immunopeptidome is stable over time and quantitative reproducible between healthy controls. **(A)** Immunopeptides identified by IMBAS-MS of a healthy donor over the course of a year. Two closely spaced time points at the start assess short term variation, and the 11-month time point is immediately post-COVID 19. Dark blue represents peptides shared between all timepoints and light blue represents peptides only measured at one time point. **(B)** Pearson correlation of immunopeptide quantities between timepoint 0 and after 12 months. **(C)** Principal component analysis of immunopeptidomes from 8 different healthy donors. Numbers refer to the different donors and colors represent the replicates. **(D)**Clustered heatmap of Jaccard similarities of immunopeptidomes between healthy donors (B). Note that 7 and 2 have only two replicates. **(E)** Unfiltered imputed Pearson correlation between healthy donors (B). The sample order was taken from the clusters built by Jaccard similarities in (C). **(F)** Filtered pairwise complete Pearson correlation of median intensities between donors showing a Jaccard similarity of more than 0.3 with at least one other donor (conditions without shared peptides are grey).

Having established temporal stability of the sHLA peptidome in a single healthy donor, we next compared the immunopeptidomes of eight healthy donors (Supplementary Table 1). A Principal Component Analysis (PCA) clearly clustered workflow replicates of the same donor but next grouped donors by shared types or supertypes (Fig 5C). Supporting this, a similar grouping emerged from pairwise Jaccard distances, which also revealed up to 50% overlap of identified peptide sequences between different donors with overlapping or similar presenting alleles (Fig. 5D).

Overall, the immunopeptidome of the different donors at best exhibited only a loose correlation (Fig 5E). However, when selecting donors with a Jaccard similarity of more than 30%, the pairwise quantitative correlation significantly improved (Fig 5F). Interestingly, donor 1 and donor 3 have a low Jaccard similarity (3%) between them - despite sharing one HLA-type (HLA-C0304); nonetheless, those 3% peptides show a high Pearson correlation.

These findings highlight the consistency and stability of the plasma immunopeptidome, further supporting its usefulness for insights into potential commonalities and variations among individuals.

## DISCUSSION AND OUTLOOK

Here we developed and applied IMBAS-MS, an improved approach to immunopeptidomics, with drastically enhanced sensitivity. This user-friendly and adaptable workflow replaces the traditional ProteinA/G affinity-based capture of anti-HLA antibodies^24^ with a streptavidin-biotin one. This enables generic use of any biotinylated antibody, regardless of their immunoglobulin type and greatly simplifies plasma-based immunopeptidomics by eliminating the need for plasma pre-clearance with its associated losses.

IMBAS-MS also eliminates nearly all hands-on time, in turn enabling the rapid preparation and acquisition of a large number of samples, which will be especially important in clinical environments. We expect IMBAS-MS to have the same advantages in tissue-based immunopeptidomics and we plan to explore this aspect in the future.

As part of our workflow, we have also implemented Data Independent Acquisition (DIA) to expand the depth of the immunopeptidomic data. To tackle the challenge of creating a suitable search space for immunopeptidomics, we employed personalized HLA peptide libraries^18^. This considerably reduces the number of potential 9mers to 12mers in a human FASTA to be searched, increasing the number of significant identifications. In contrast to other library generation strategies^25,26^, our approach eliminates the need for any upfront measurements and can be transferred between MS platforms. It also avoids building a library from Data Dependent Acquisition (DDA) runs and could be adapted to supertype or study-specific libraries, potentially incorporating common post-translational modifications.

As a next step, we envision combining IMBAS-MS with multiplexed DIA and in particular to use one of the channels as a reference channel^27,28^. By decoupling identification and quantification, the reference channel improves proteomics depth, sensitivity and comparability between samples.

Our results highlight the potential diagnostic applicability beyond identifying cancer neoepitopes. They demonstrate the presence of very large numbers of immunopeptides in plasma samples, further supporting the notion of plasma as a valuable, non-invasive source of immunopeptides^4,11,29^.

We observed that the immunopeptides found in plasma are mostly representative of the tissue proteome. However, brain-associated proteins where less represented and it would be interesting to investigate mechanisms of presentation of these sHLAs in the plasma. We also demonstrated the existence of a stable healthy plasma immunopeptidome, both quantitatively and qualitatively, across different healthy individuals. This finding is highly relevant for clinical applications, as it suggests that a general baseline healthy immunopeptidome can be established. In turn, this could significantly facilitate the identification of disease-specific immunopeptide signatures and aid in the development of novel diagnostic markers and therapeutic strategies. Such an approach could extend the diagnostic potential of plasma immunopeptidome profiling within and beyond the search for neoepitopes in the context of cancer. This may provide insights into a wide range of pathological conditions that involve alterations in immune responses, such as autoimmune disorders, infectious diseases or inflammatory conditions. In this context, the minimal-invasive nature of plasma-based immunopeptidome profiling combined with the streamlined IMBAS-MS technology could enable a patient-friendly approach to disease monitoring and personalized medicine, facilitating earlier intervention and more effective treatment strategies.

Clearly, future studies are needed to expand upon these exciting findings by investigating basic aspects of sHLA generation and presentation and the diagnostic capabilities of plasma immunopeptide signatures in specific disease states. Combined with ongoing development of the underlying analytical technology, sHLA peptidomics may become an important addition to the arsenal of precision medicine.

## ABBREVIATIONS

ACN: acetonitrile
CAR: T-cell chimeric antigen receptor T-cell
CTL: cytotoxic T lymphocytes
CYP: Cytochrome-P-Oxygenase
dda: data-dependent acquisition
dia: data-independent acquisition
FA: formic acid
GO: Gene Ontology
(s/m)HLA: (soluble/membrane) human leukocyte antigen
IgG: Immunoglobuline G
IEDB: Immune Epitope Database & Tools
IM: ion mobility
IMBAS: Streptavidin based immunopeptidomics workflow (IMmunopeptidomics by Biotinylated Antibodies and Streptavidin)
MeOH: methanol
MHC: major histocompatibility complex
PASEF: parallel accumulation – serial fragmentation
PBS: phosphate-buffered saline
py_diAID: Python package for Data-Independent Acquisition with an Automated Isolation Design
SPD: samples per day
TIMS: trapped ion mobility spectrometry

## ACKNOWLEDGEMENTS

This study was supported by the Max Planck Society for Advancement of Science, the European Union’s Horizon 2020 research and innovation program under grant agreement No 874839 (ISLET) and by the Bavarian State Ministry of Health and Care through the research project DigiMed Bayern (www.digimed-bayern.de). MW, MT and MZ acknowledge support from the International Max Planck Research School for Life Sciences – IMPRS-LS. We are grateful for the sequencing support provided by the NGS Core Facility at the Max Planck Institute of Biochemistry. We are especially grateful for the blood donors. We thank our colleagues in the Department of Proteomics and Signal Transduction, Max Planck Institute of Biochemistry and at the Center for Protein Research at Copenhagen University, for discussions and support. In particular, we thank I. Paron and T. Heymann for technical support and Medini Steger for scientific administration support.

## AUTHOR CONTRIBUTIONS

MW, MT and MM conceptualized and designed the study. MW designed and performed experiments. W-FZ predicted AlphaPeptDeep libraries. MW and MZ transferred the workflow to the automation platform. MZ performed the HLA-typing. MW and MT analyzed the data. MW and PS designed the MS acquisition methods. MW and MM wrote the original manuscript draft. All authors read, revised and approved the manuscript.

## DATA AVAILABILITY

The raw mass spectrometry data have been deposited in the public proteomics repository and will be available after publication.

## SUPPLEMENTAL DATA

This article contains supplemental data.

