## Supplements for "The potential of plasma HLA peptides beyond neoepitopes"

**Supplementary Table 1:** Donor identifier with associated HLA-type including Figure assignments.

| Donor | HLA-type | Figure 1 | Figure 2 | Figure 3 | Figure 5 |
|---|---|---|---|---|---|
| 1 | HLA-A24:02,HLA-A26:01,HLA-B40:01,HLA-B39:01,HLA-C03:04,HLA-C12:03 | x | x | x (Donor 1) | x (Donor 1) |
| 2 | HLA-A24:02, HLA-A31:01, HLA-B27:07, HLA-B50:01, HLA-C06:02, HLA-C15:02 | - | - | x (Donor 2) | x (Donor 2) |
| 3 | HLA-A02:01, HLA-A32:01, HLA-B15:01, HLA-B35:01, HLA-C03:04, HLA-C04:01 | - | - | x (Donor 3) | x (Donor 3) |
| 4 | HLA-A02:01,HLA-A32:01,HLA-B08:01,HLA-B35:03,HLA-C04:01,HLA-C07:01 | - | - | x (Donor 5) | x (Donor 4);timeline* |
| 5 | HLA-A02:06, HLA-A11:01, HLA-B15:13, HLA-B40:01, HLA-C08:01, HLA-C15:02 | - | - | - | x (Donor 5) |
| 6 | HLA-A01:01, HLA-A25:01, HLA-B07:02, HLA-B55:01, HLA-C03:03, HLA-C07:02 | - | - | - | x (Donor 6) |
| 7 | HLA-A24:02, HLA-A30:01, HLA-B07:43, HLA-B07:236, HLA-C07:02, HLA-C08:02 | - | - | x (Donor 6) | x (Donor 7) |
| 8 | HLA-A01:01, HLA-A24:03,HLA-B38:20, HLA-B57:01, HLA-C06:02, HLA-C12:03 | - | - | x (Donor 4) | x (Donor 8) |

* 11 month sampling point post mild COVID-19 infection (confirmed by antigen test) was taken 3 days after first negative antigen test.

A)

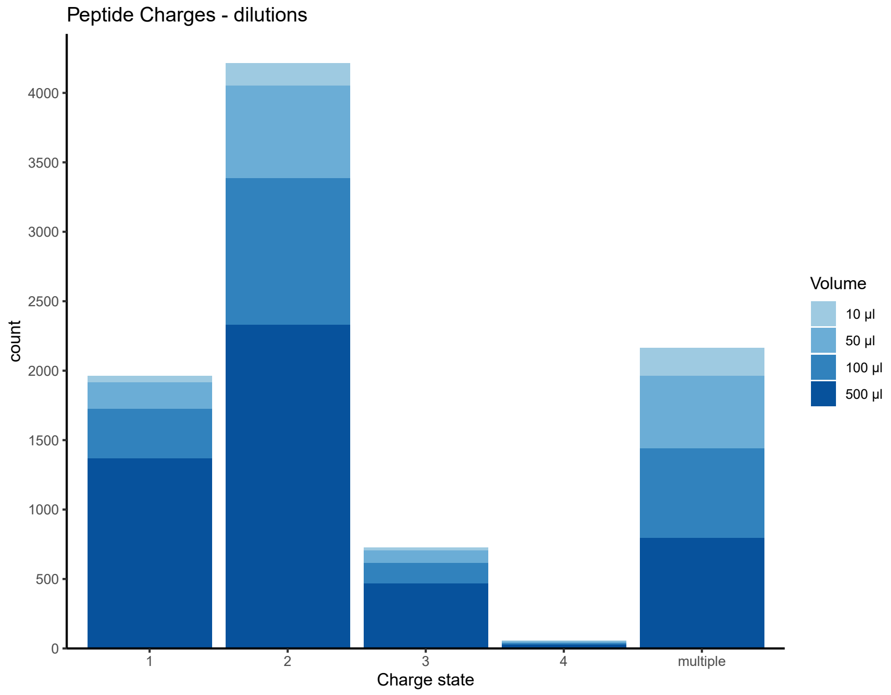
Peptide Charges - dilutions
count
4000
3500
3000
2500
2000
1500
1000
500
0
1
2
3
4
multiple
Charge state
Volume
10 µl
50 µl
100 µl
500 µl

B)

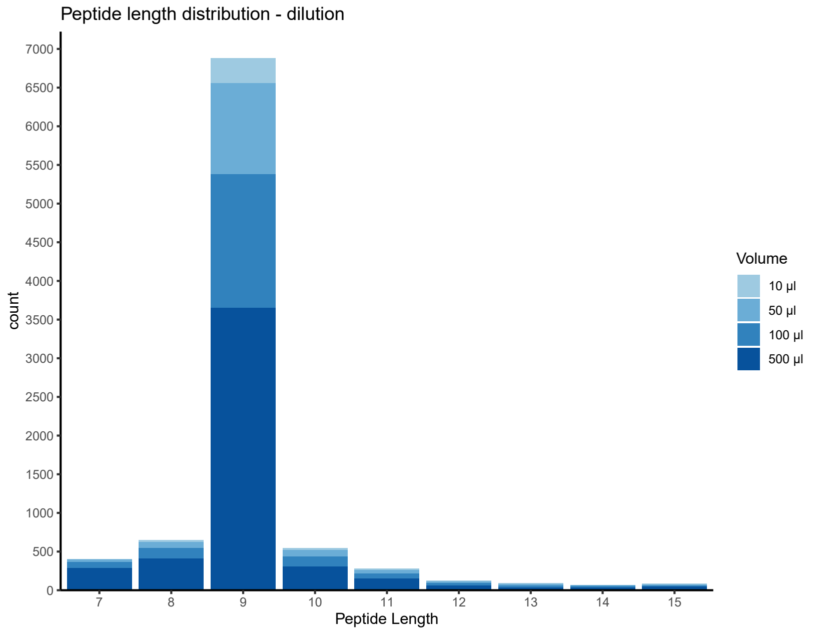
Peptide length distribution - dilution
count
7000
6500
6000
5500
5000
4500
4000
3500
3000
2500
2000
1500
1000
500
0
7
8
9
10
11
12
13
14
15
Peptide Length
Volume
10 µl
50 µl
100 µl
500 µl

**Supplementary Figure 1: Length and charge properties of IMBAS-MS DDA based identifications. (A)** Stacked bar plot of charge states of all identified precursors identified in the dilution series experiment (Figure 1C). **(B)** Stacked bar plot of length distribution of identified HLA class I peptides of all samples in (Figure 1C).

A)

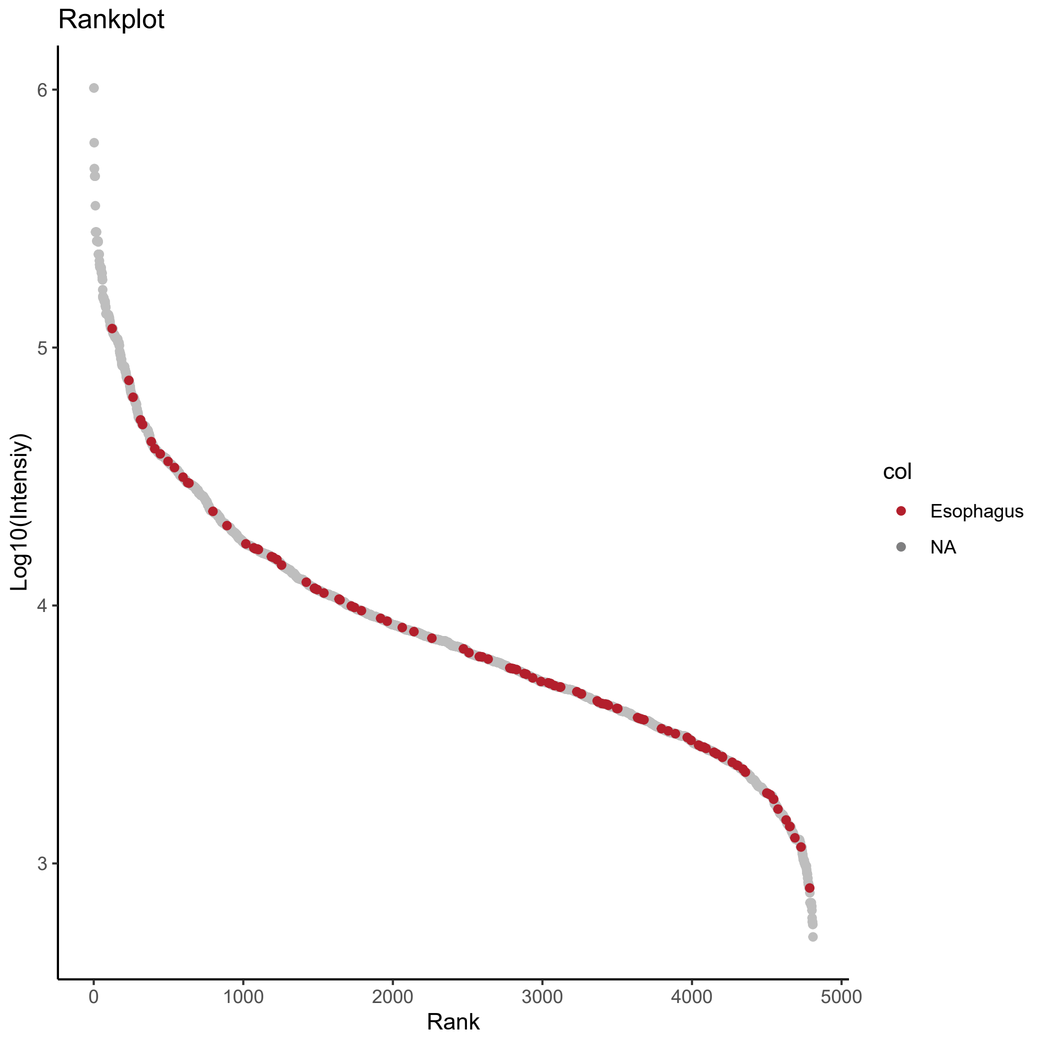
Rankplot
Log10(Intensiy)
6
5
4
3
0
1000
2000
3000
4000
5000
Rank
col
Esophagus
NA

B)

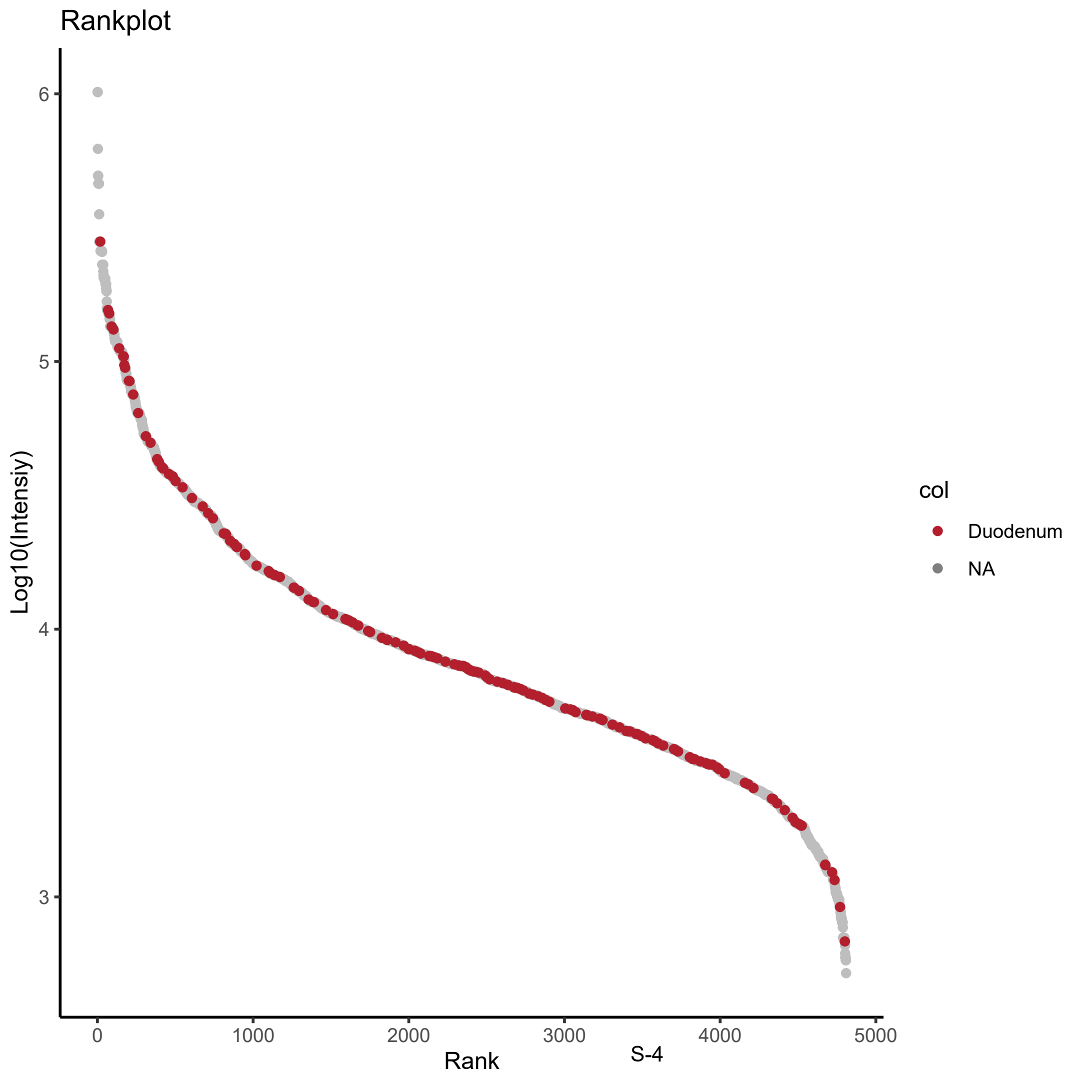
Rankplot
Log10(Intensiy)
6
5
4
3
0
1000
2000
3000
4000
5000
Rank
col
Duodenum
NA

C)

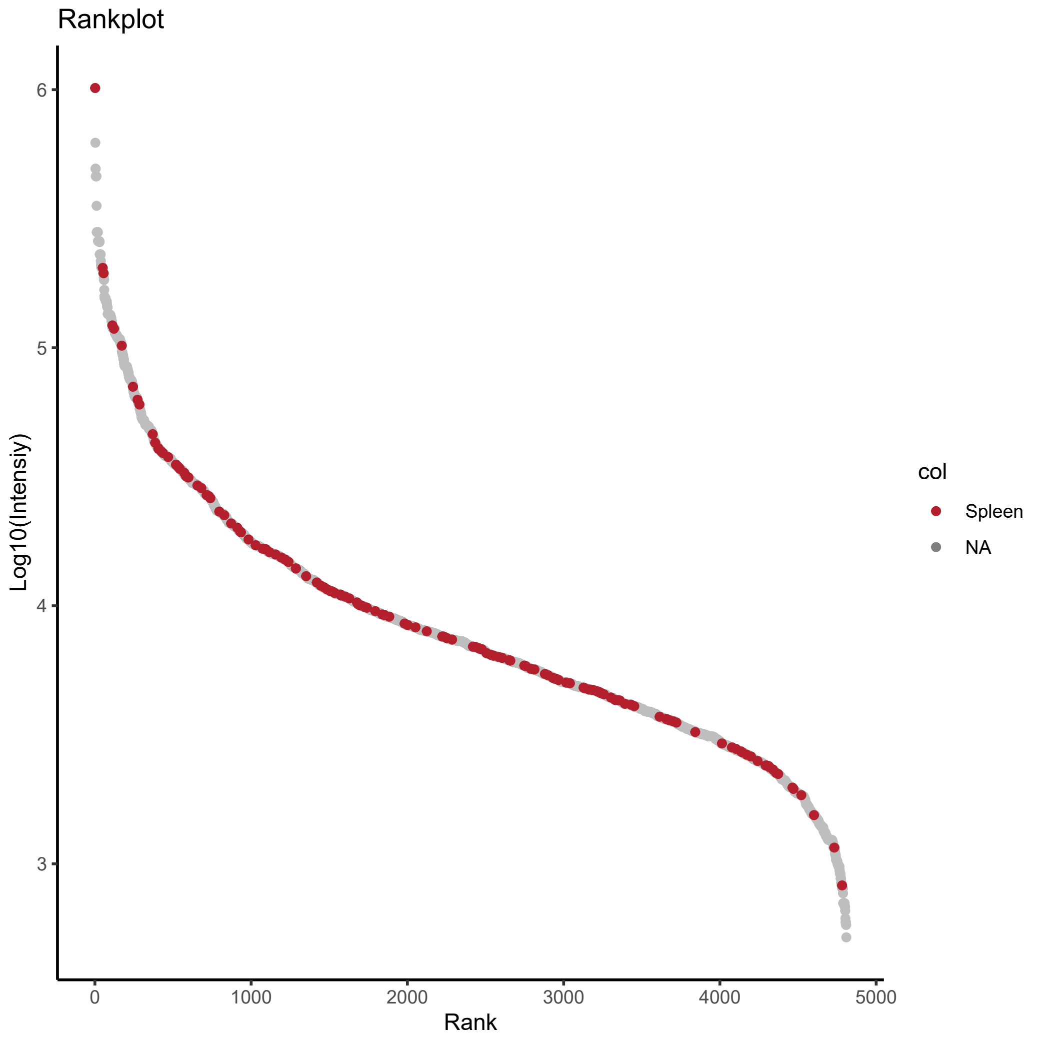

Rankplot
Log10(Intensiy)
6
5
4
3
0
1000
2000
3000
4000
5000
Rank
col
Spleen
NA


D)

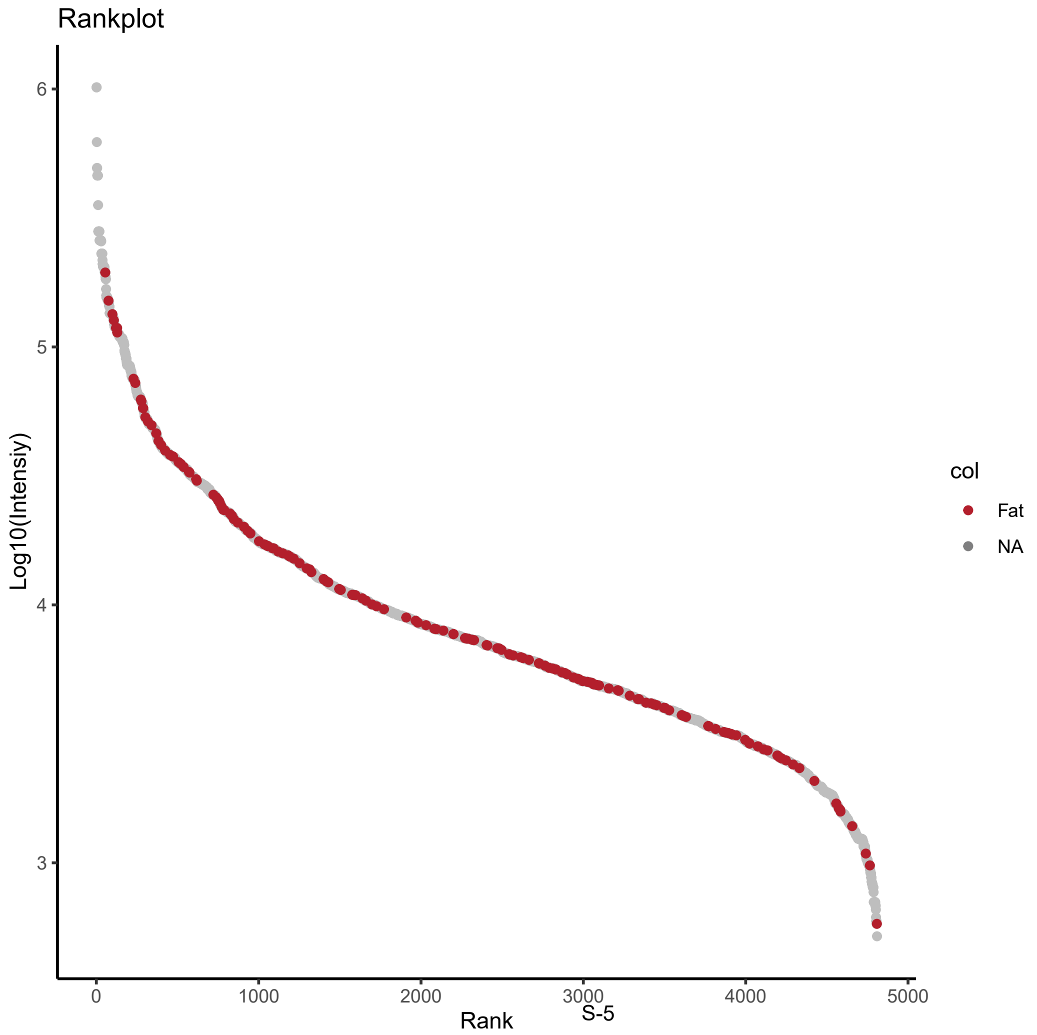

Rankplot
Log10(Intensiy)
6
5
4
3
0
1000
2000
3000
4000
5000
Rank
col
Fat
NA

E)

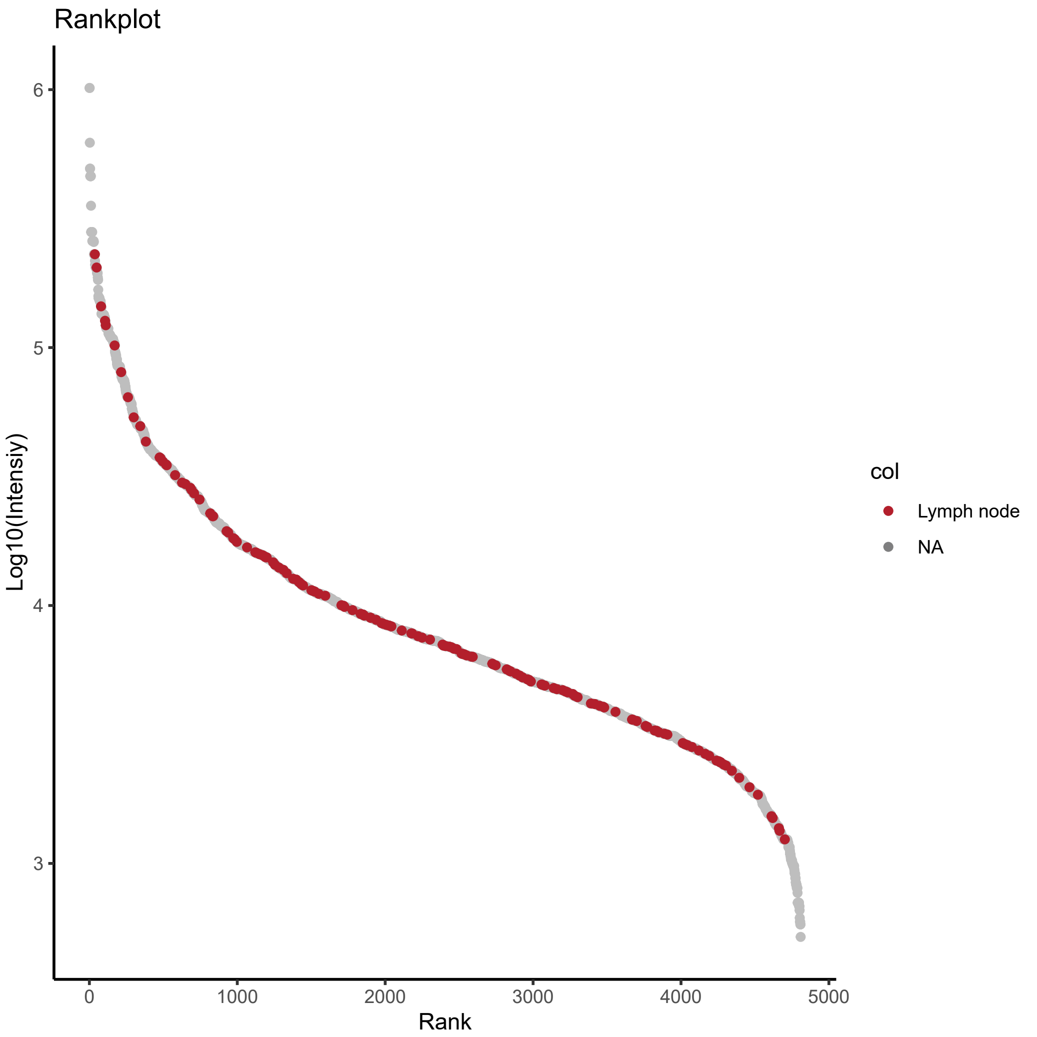
Rankplot
Log10(Intensiy)
6
5
4
3
0
1000
2000
3000
4000
5000
Rank
col
Lymph node
NA

F)

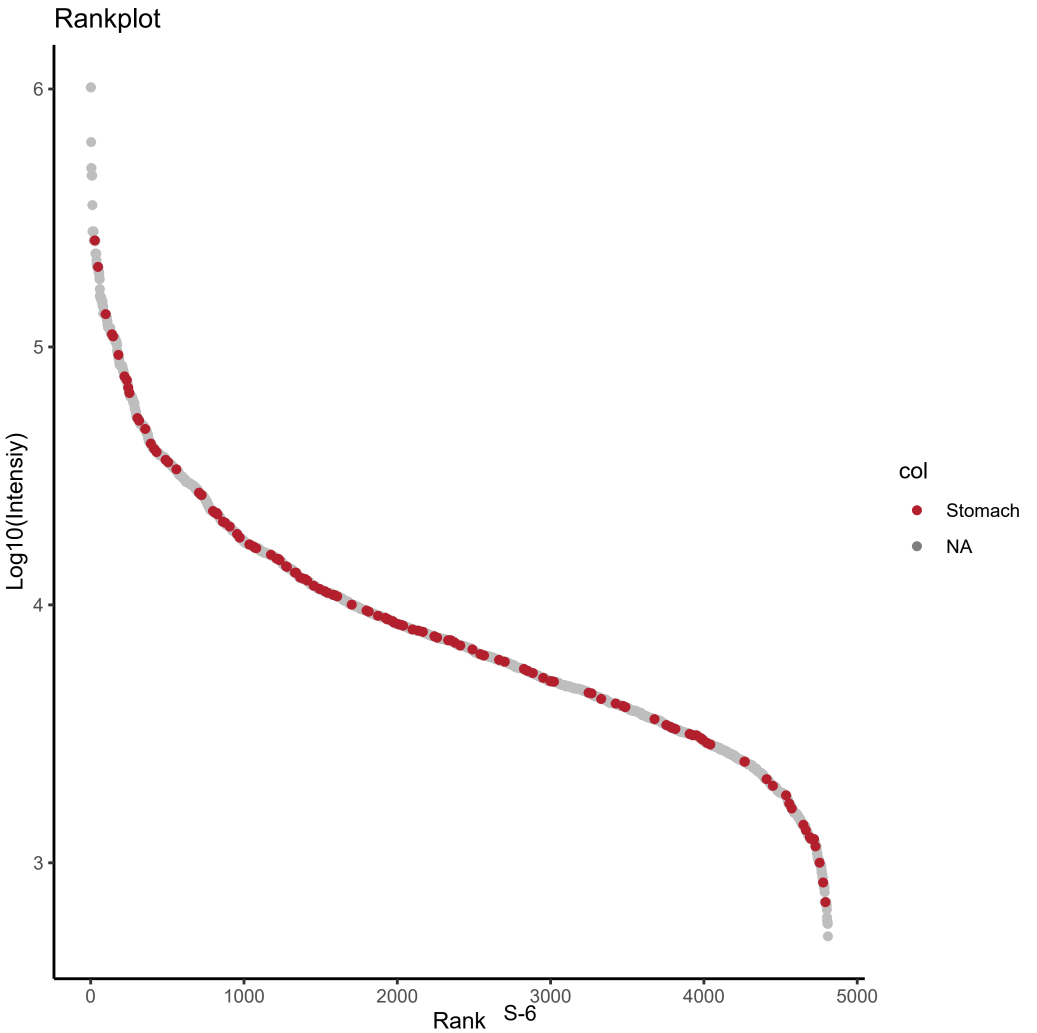
Rankplot
Log10(Intensiy)
6
5
4
3
0
1000
2000
3000
4000
5000
Rank
col
Stomach
NA

I)

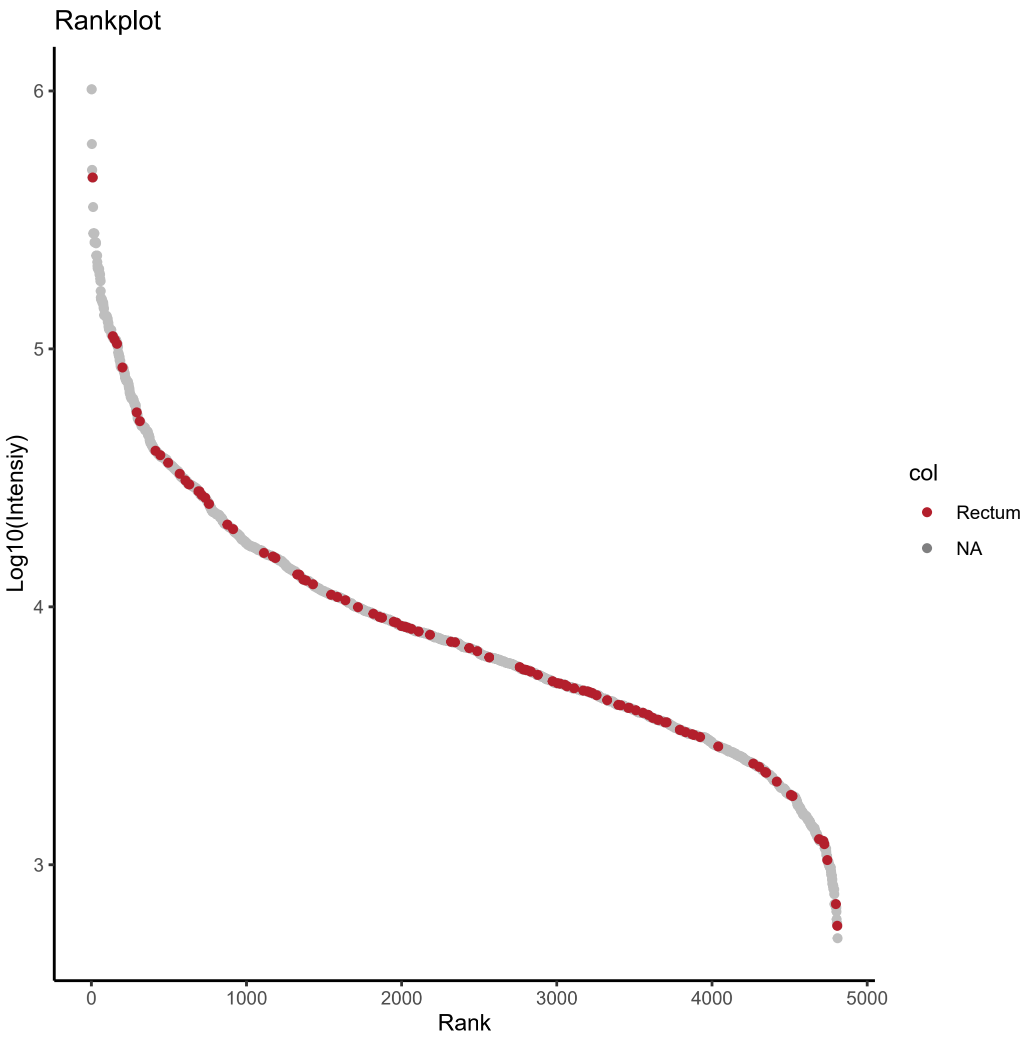
Rankplot
6
5
4
3
Log10(Intensiy)
0
1000
2000
3000
4000
5000
Rank
col
Rectum
NA

J)

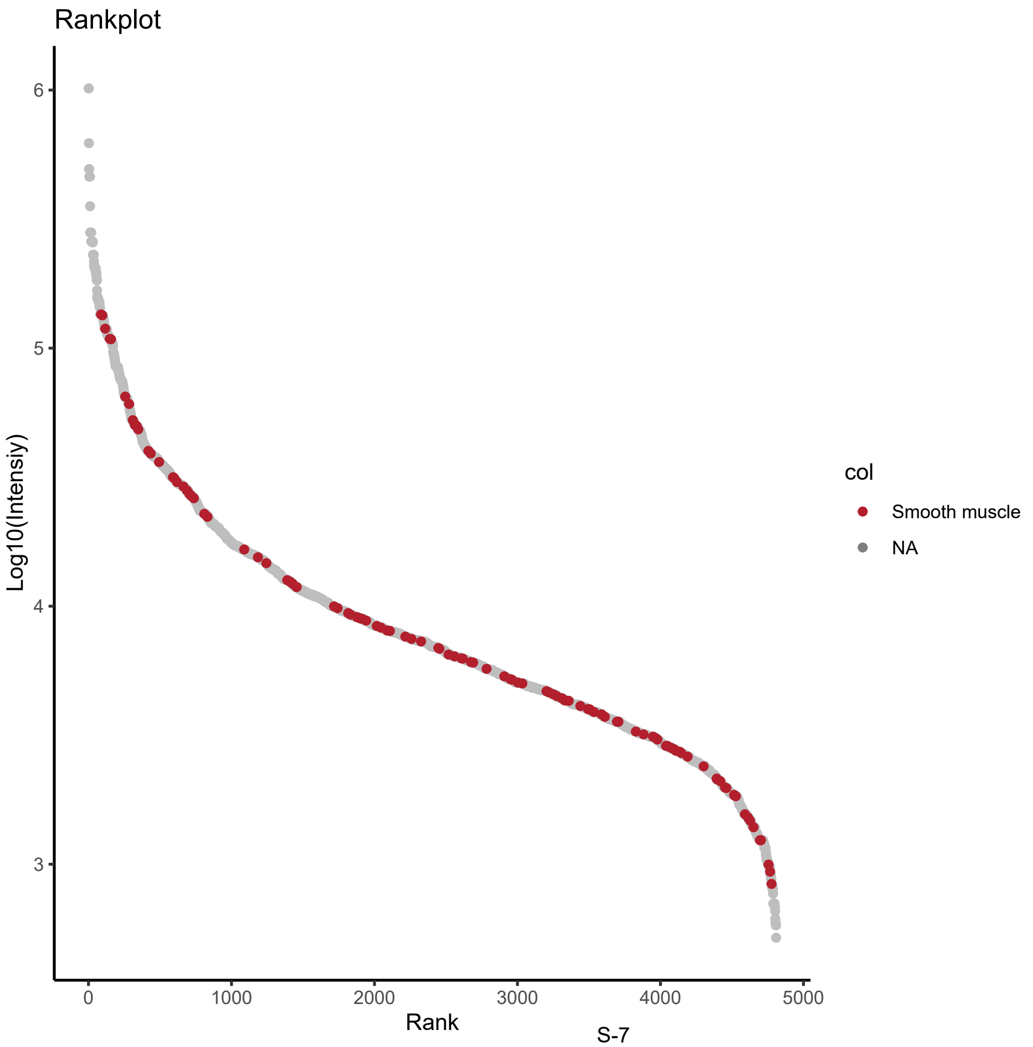
Rankplot
6
5
4
3
Log10(Intensiy)
0
1000
2000
3000
4000
5000
Rank
col
Smooth muscle
NA

K)

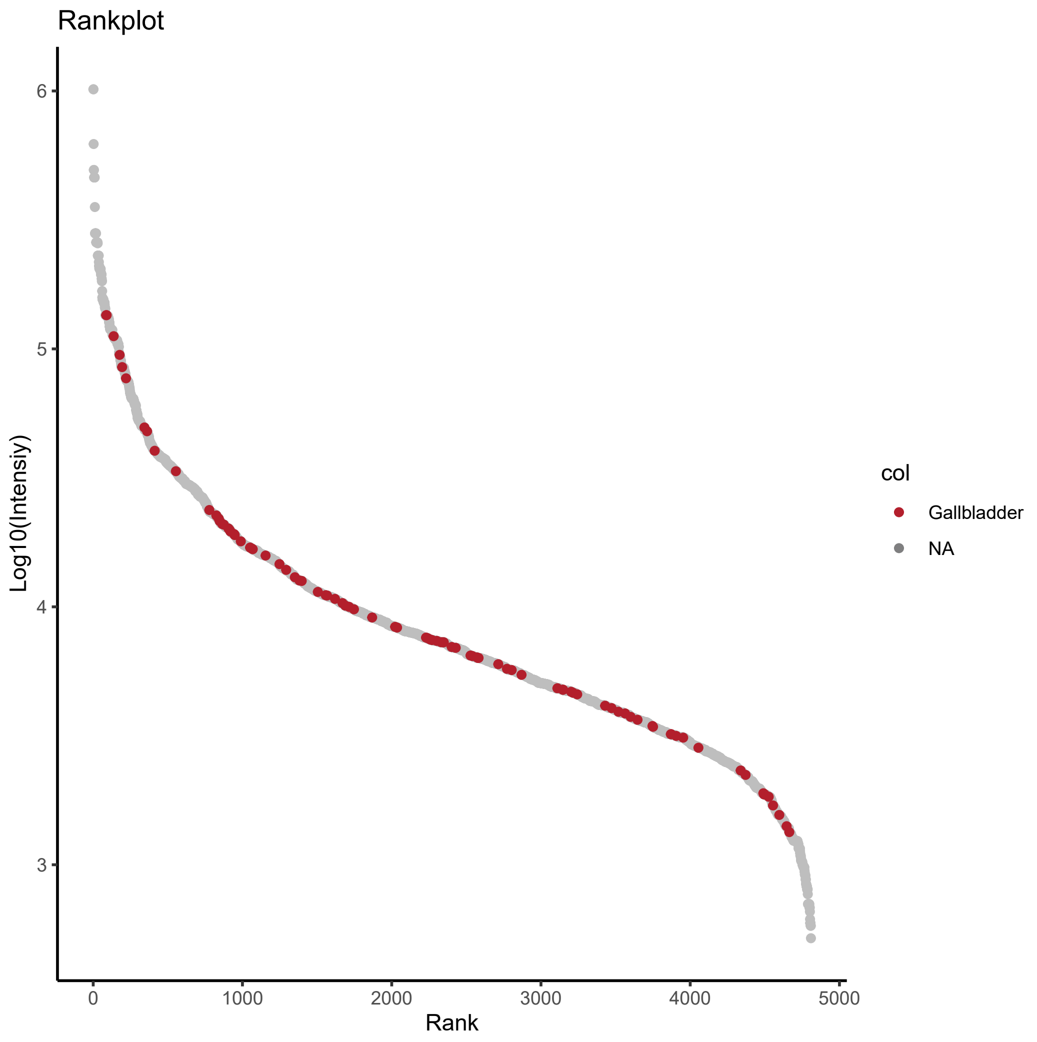
Rankplot
Log10(Intensiy)
6
5
4
3
0
1000
2000
3000
4000
5000
Rank
col
Gallbladder
NA

L)

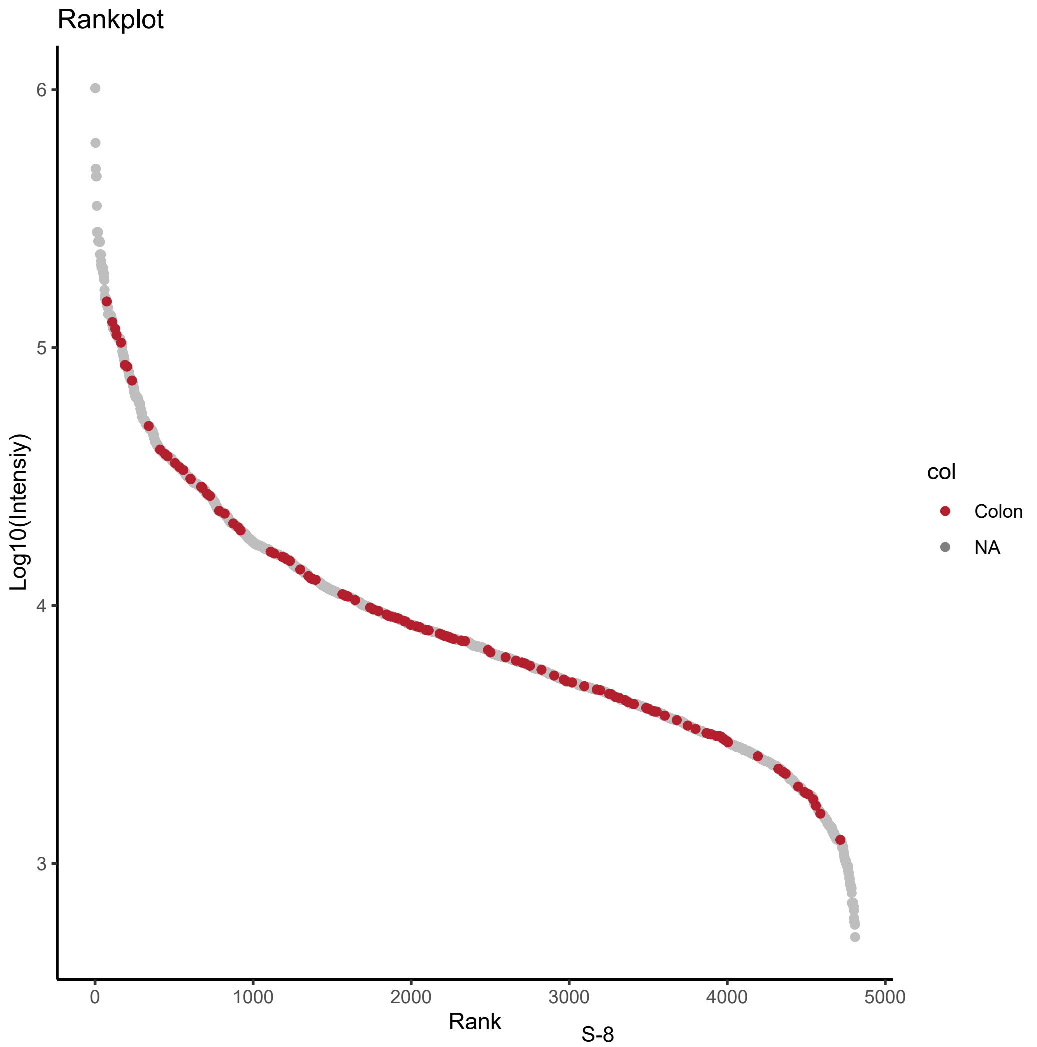
Rankplot
Log10(Intensiy)
6
5
4
3
0
1000
2000
3000
4000
5000
Rank
col
Colon
NA

M)

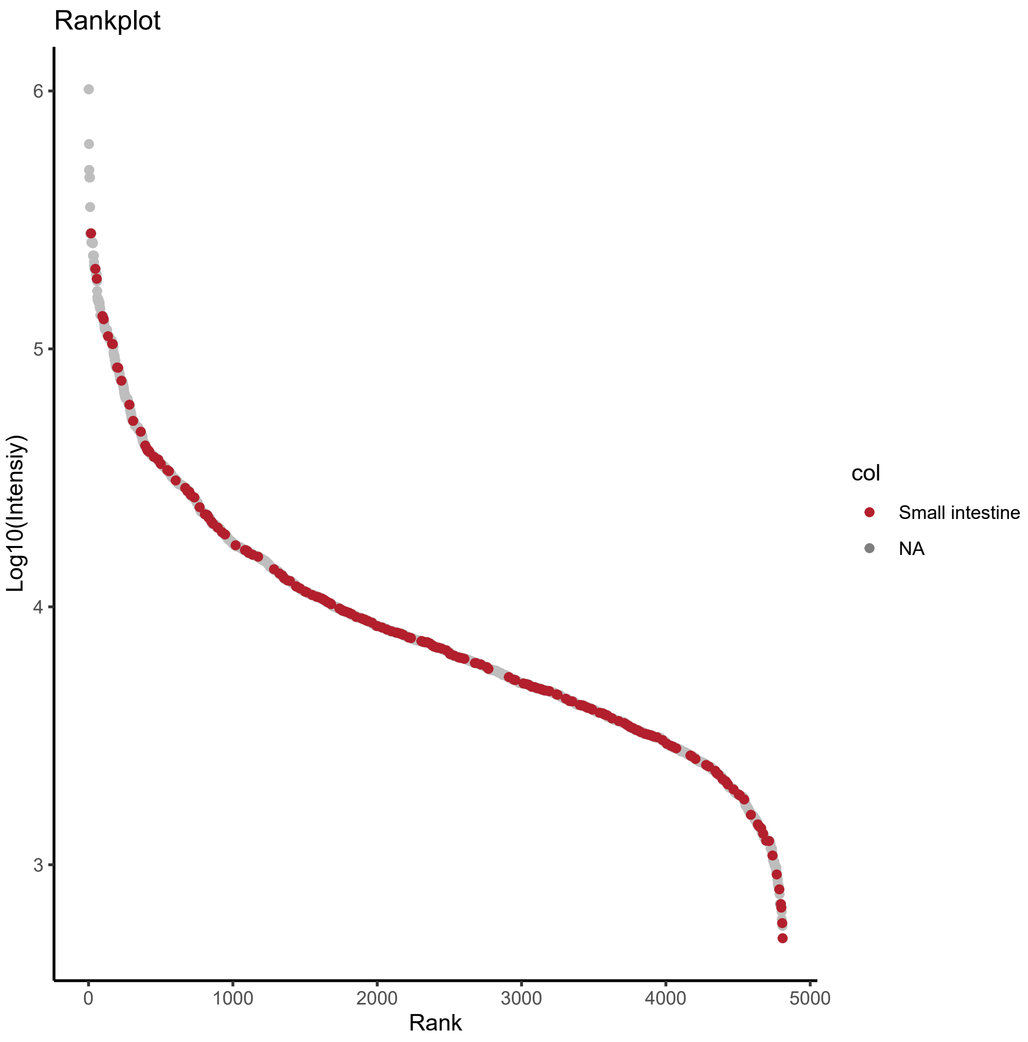
Rankplot
Log10(Intensiy)
6
5
4
3
0
1000
2000
3000
4000
5000
Rank
col
Small intestine
NA

N)

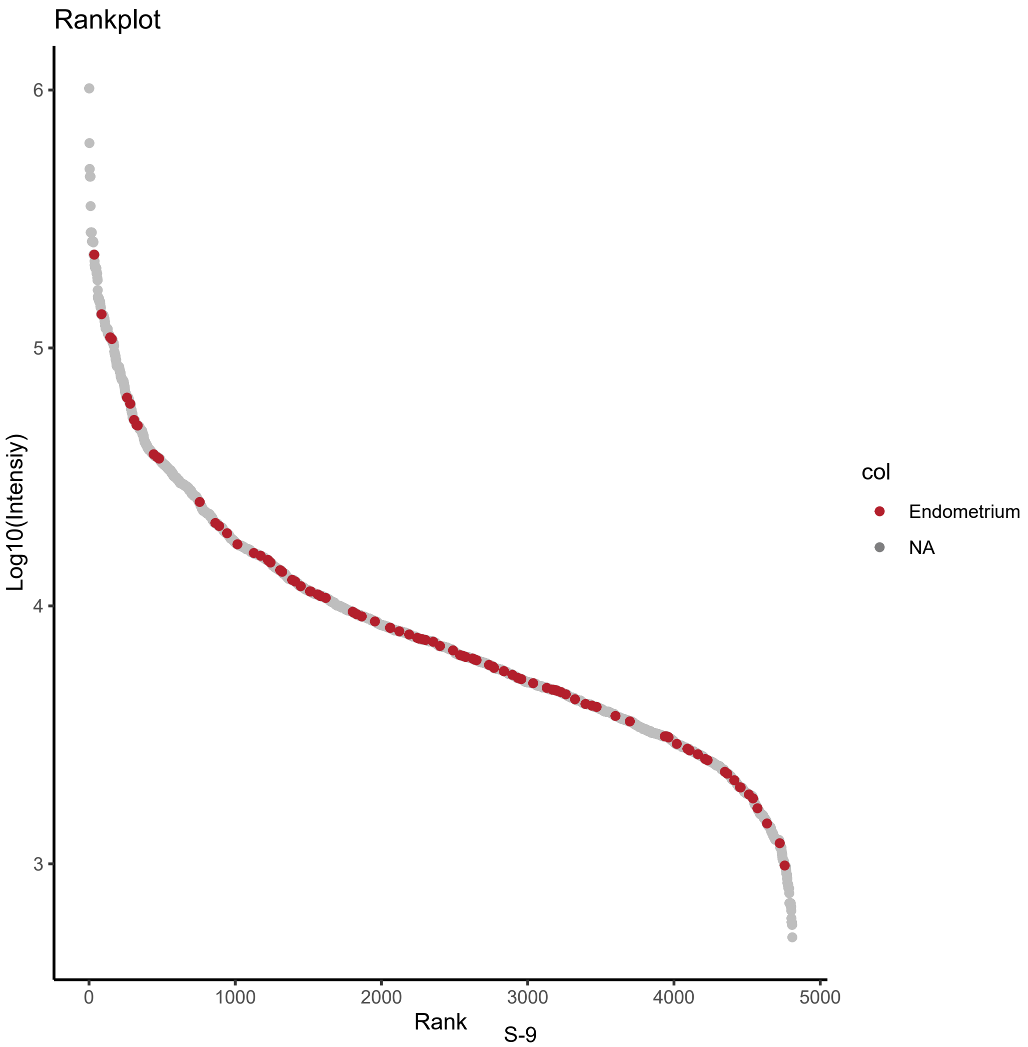
Rankplot
Log10(Intensiy)
6
5
4
3
0
1000
2000
3000
4000
5000
Rank
col
Endometrium
NA

G)

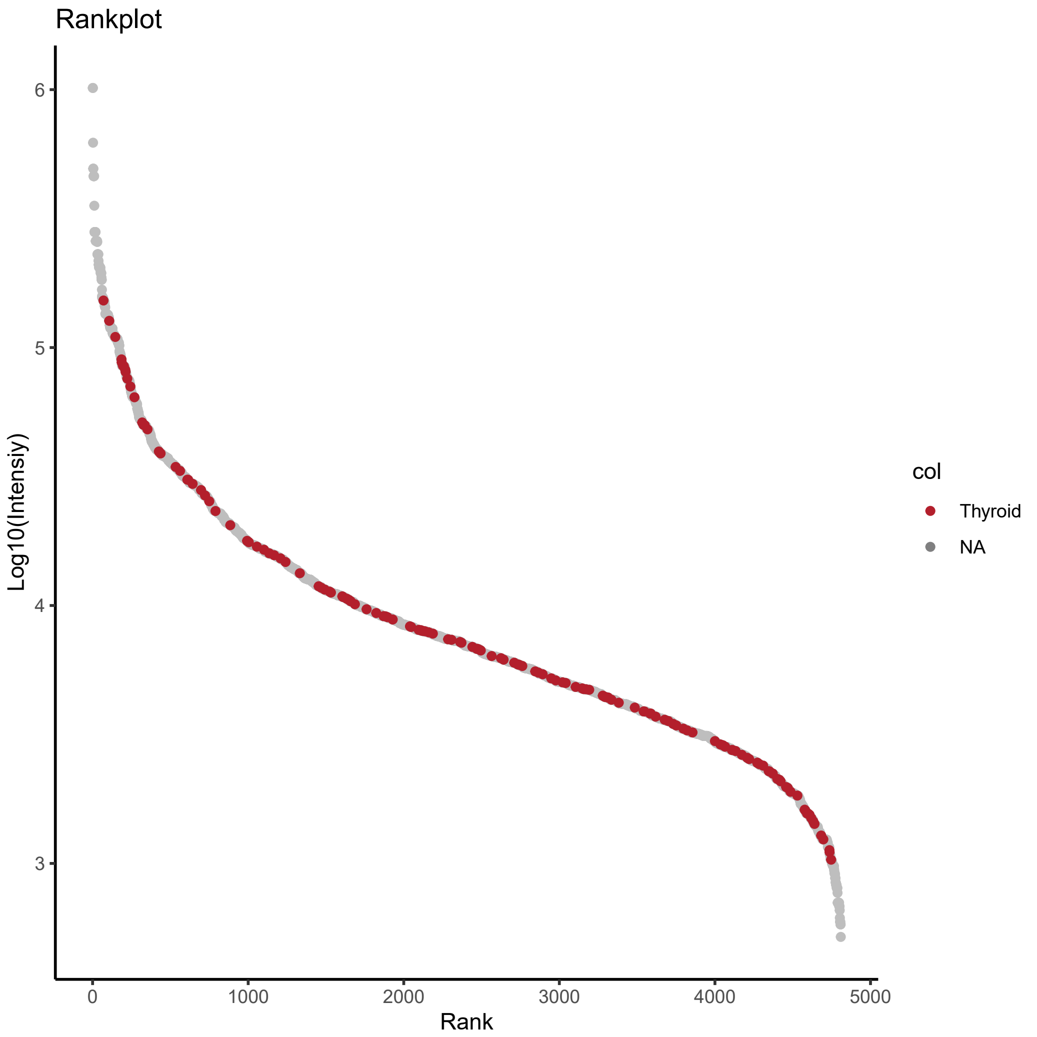
Rankplot
Log10(Intensiy)
6
5
4
3
0
1000
2000
3000
4000
5000
Rank
col
Thyroid
NA

H)

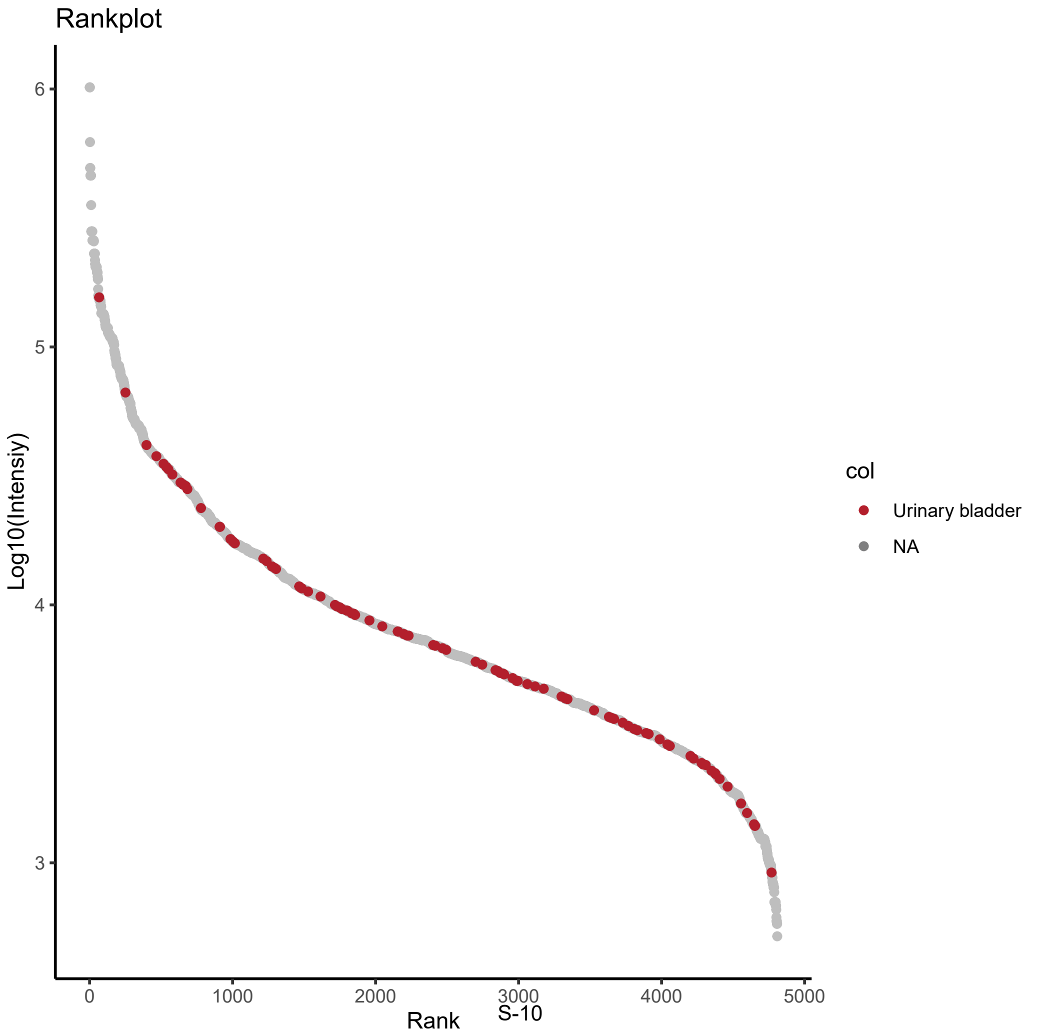
Rankplot
Log10(Intensiy)
6
5
4
3
0
1000
2000
3000
4000
5000
Rank
col
Urinary bladder
NA

O)

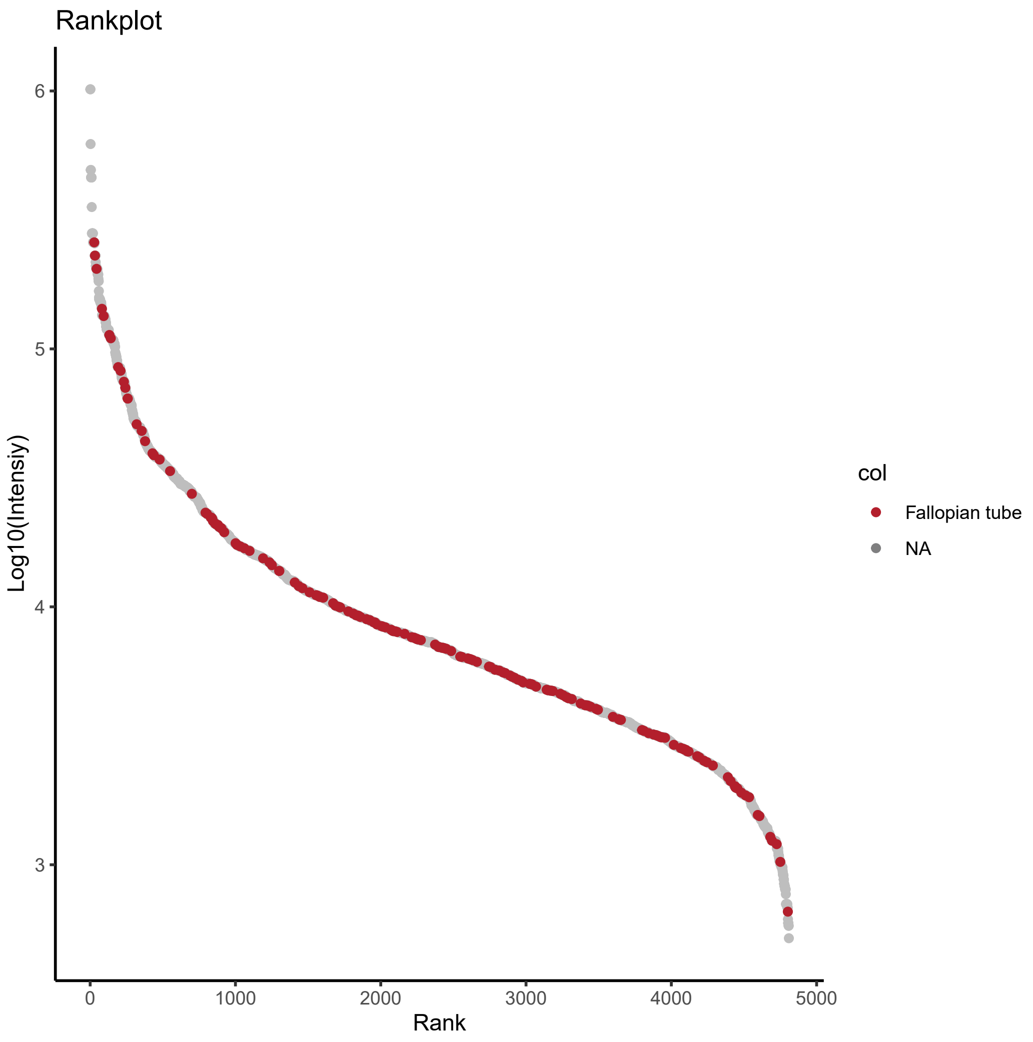
Rankplot
Log10(Intensiy)
6
5
4
3
0
1000
2000
3000
4000
5000
Rank
col
Fallopian tube
NA

P)

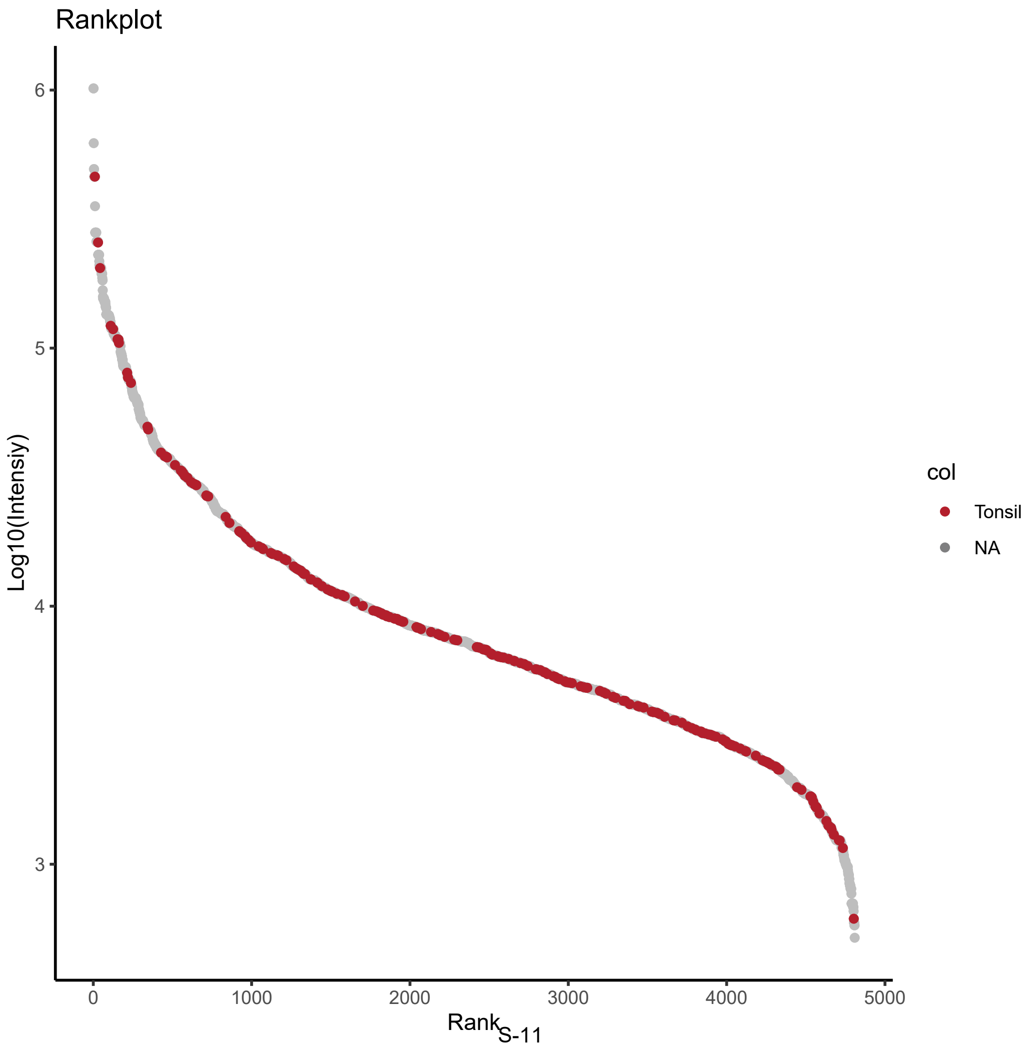
Rankplot
Log10(Intensiy)
6
5
4
3
0
1000
2000
3000
4000
5000
Rank
col
Tonsil
NA

Q)

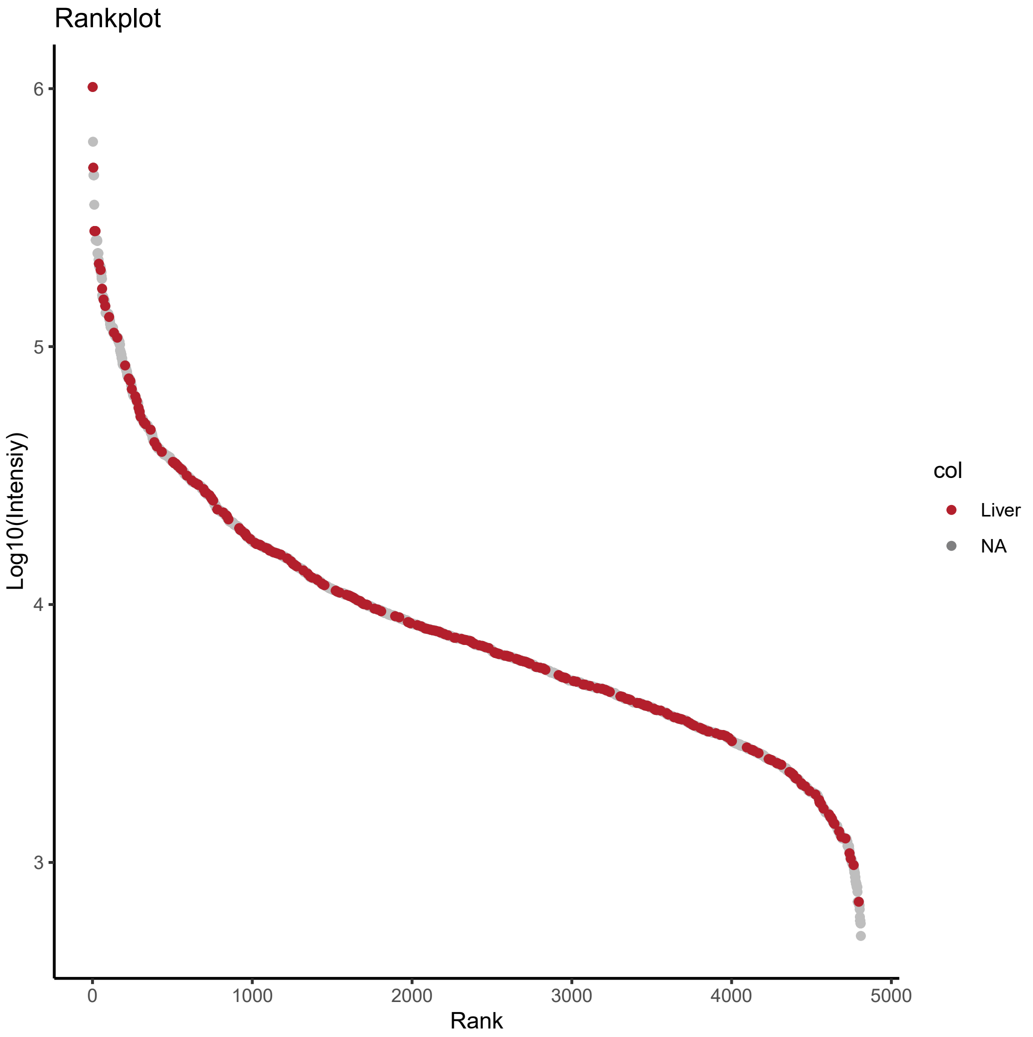
Rankplot
6
5
4
3
Log10(Intensiy)
0
1000
2000
3000
4000
5000
Rank
col
Liver
NA

R)

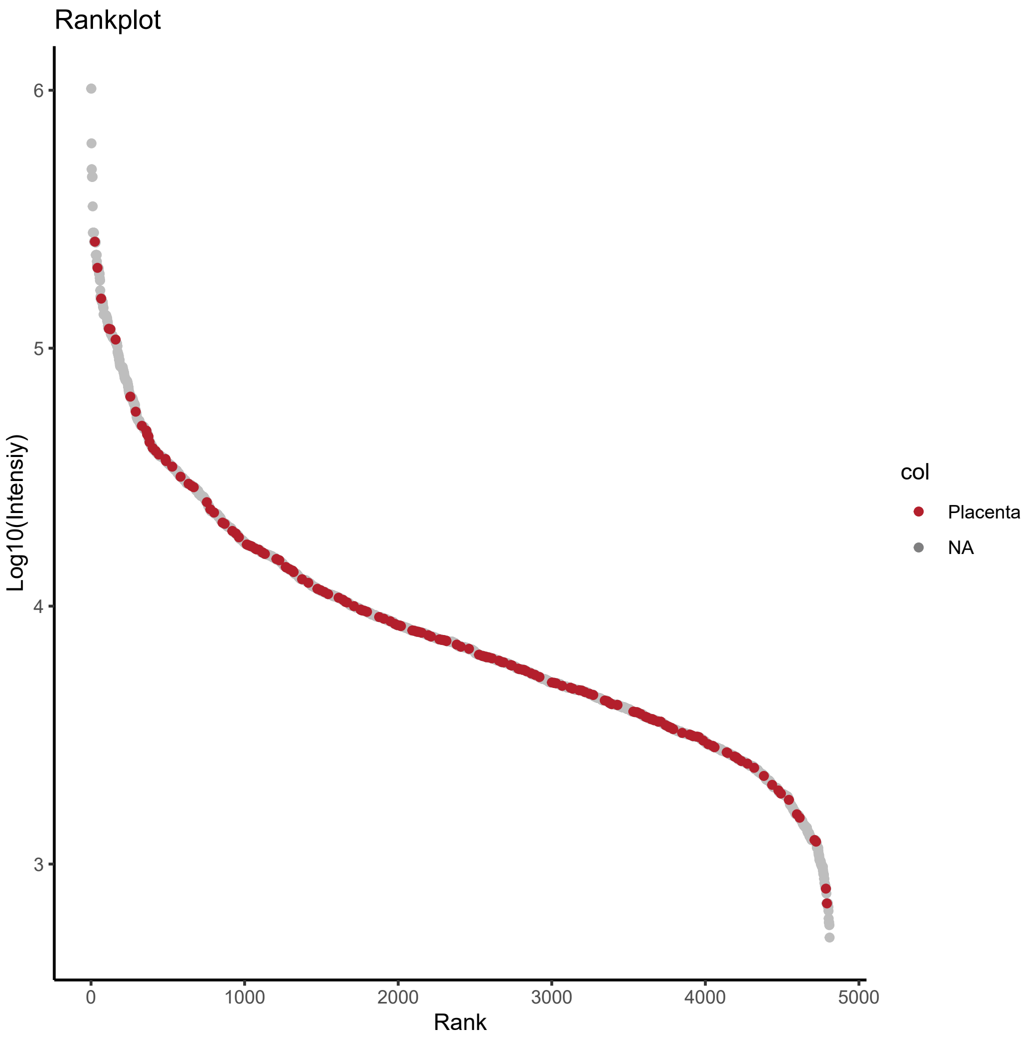
Rankplot
6
5
4
3
Log10(Intensiy)
0
1000
2000
3000
4000
5000
Rank
col
Placenta
NA

S)

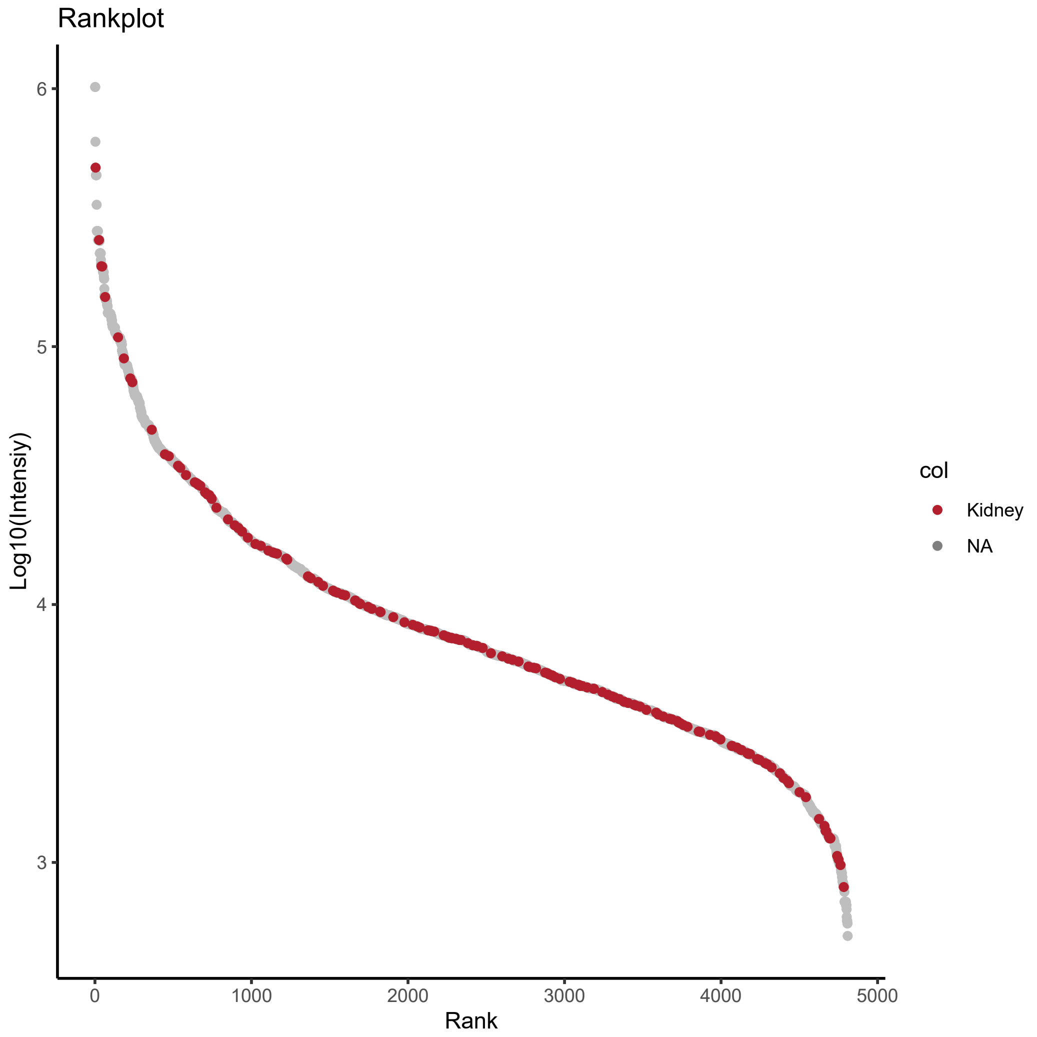
Rankplot
Log10(Intensiy)
6
5
4
3
0
1000
2000
3000
4000
5000
Rank
col
Kidney
NA

T)

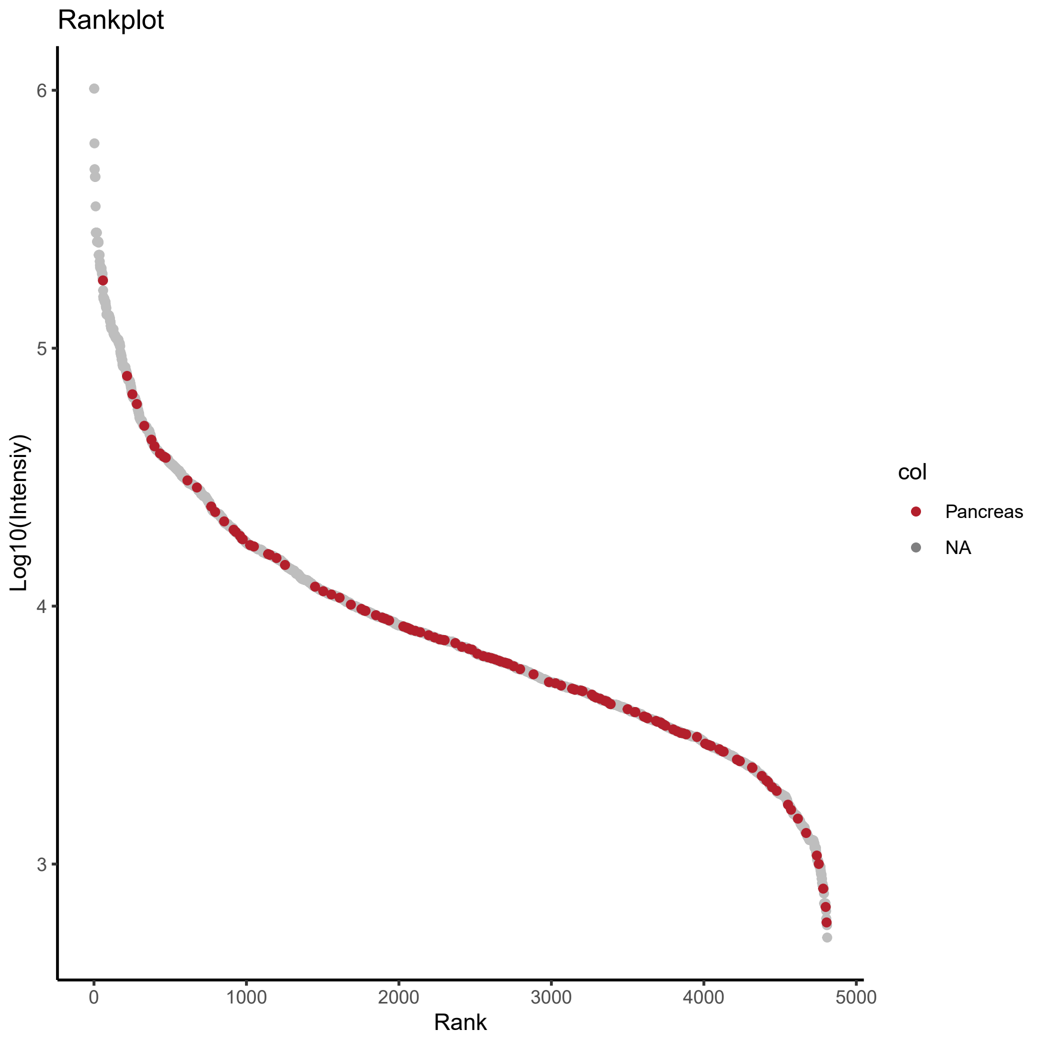
Rankplot
Log10(Intensiy)
6
5
4
3
0
1000
2000
3000
4000
5000
Rank
col
Pancreas
NA

U)

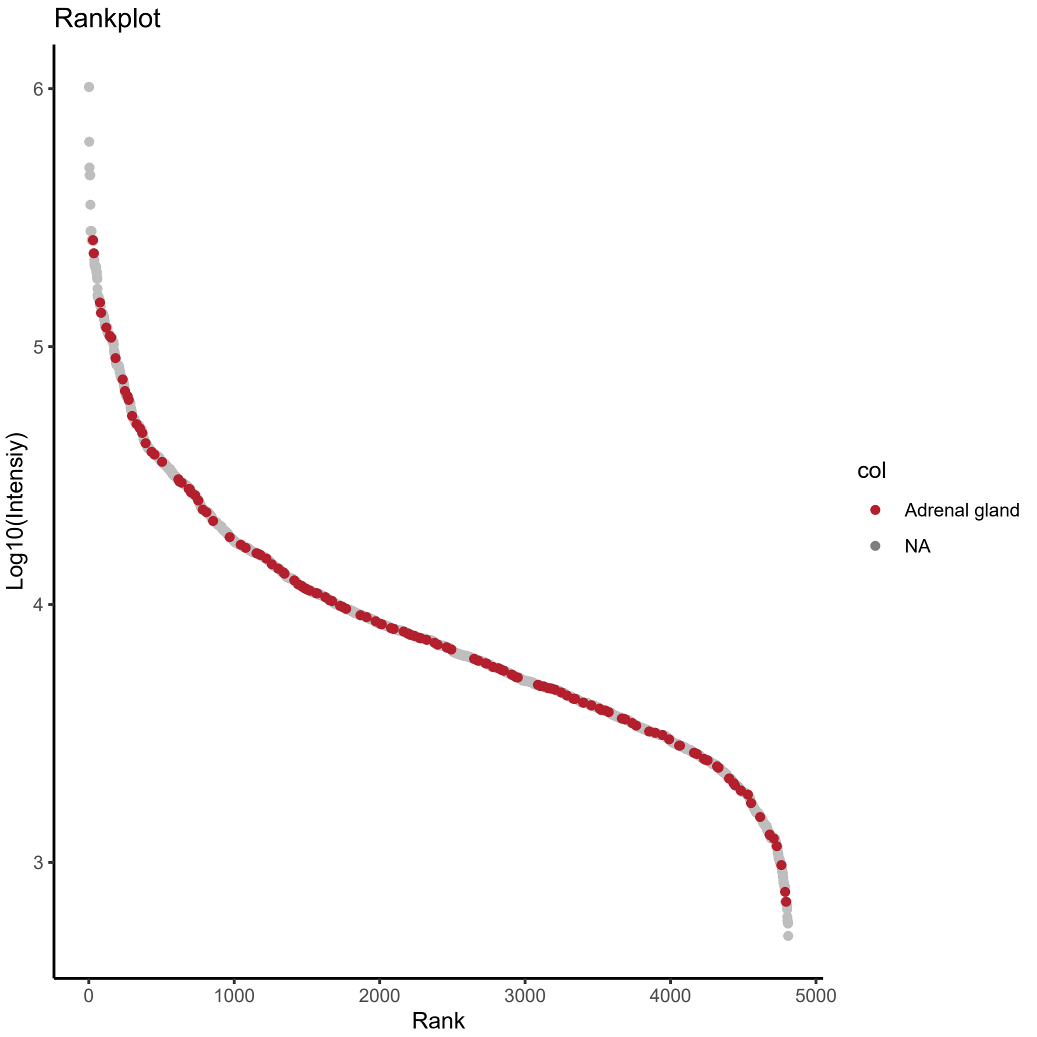
Rankplot
Log10(Intensiy)
6
5
4
3
0
1000
2000
3000
4000
5000
Rank
col
Adrenal gland
NA

V)

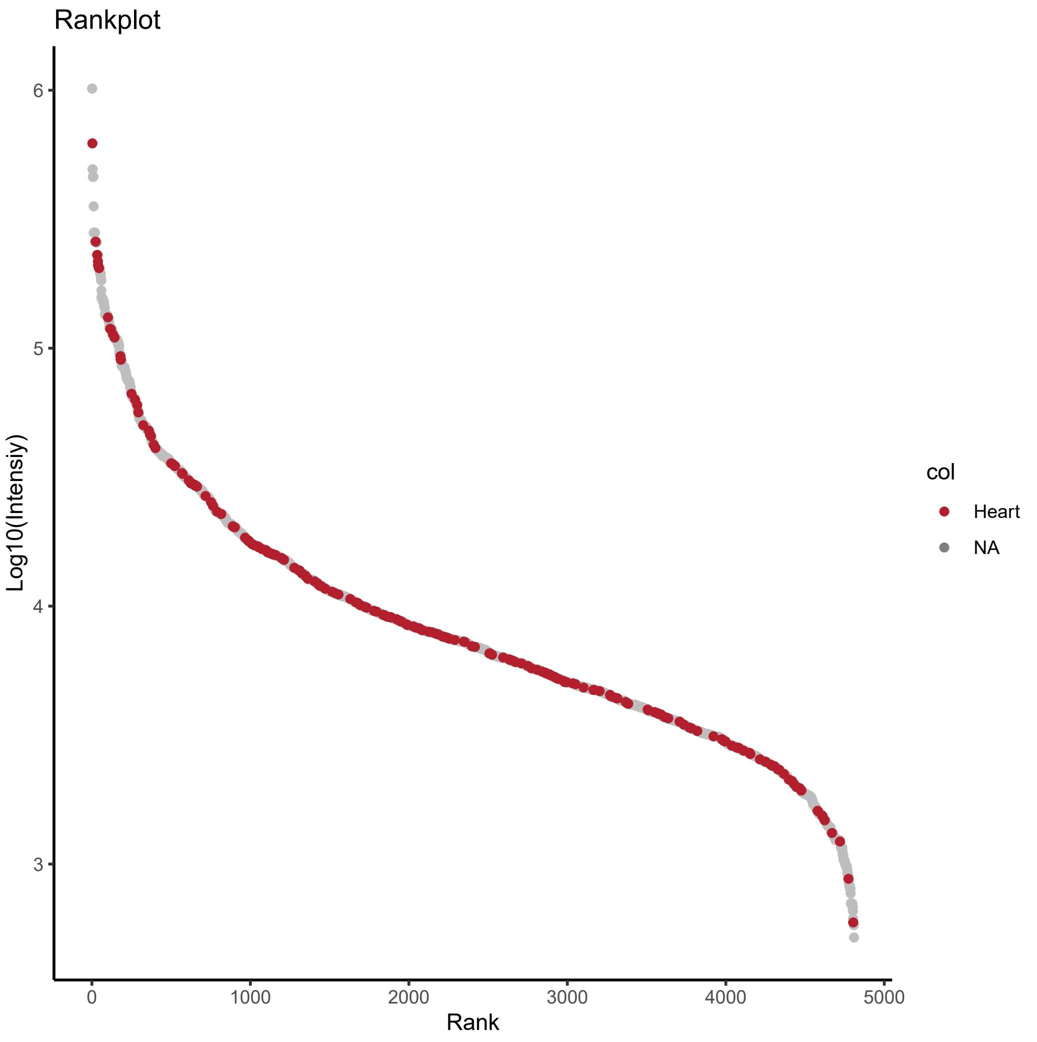
Rankplot
Log10(Intensiy)
6
5
4
3
0
1000
2000
3000
4000
5000
Rank
col
Heart
NA

W)

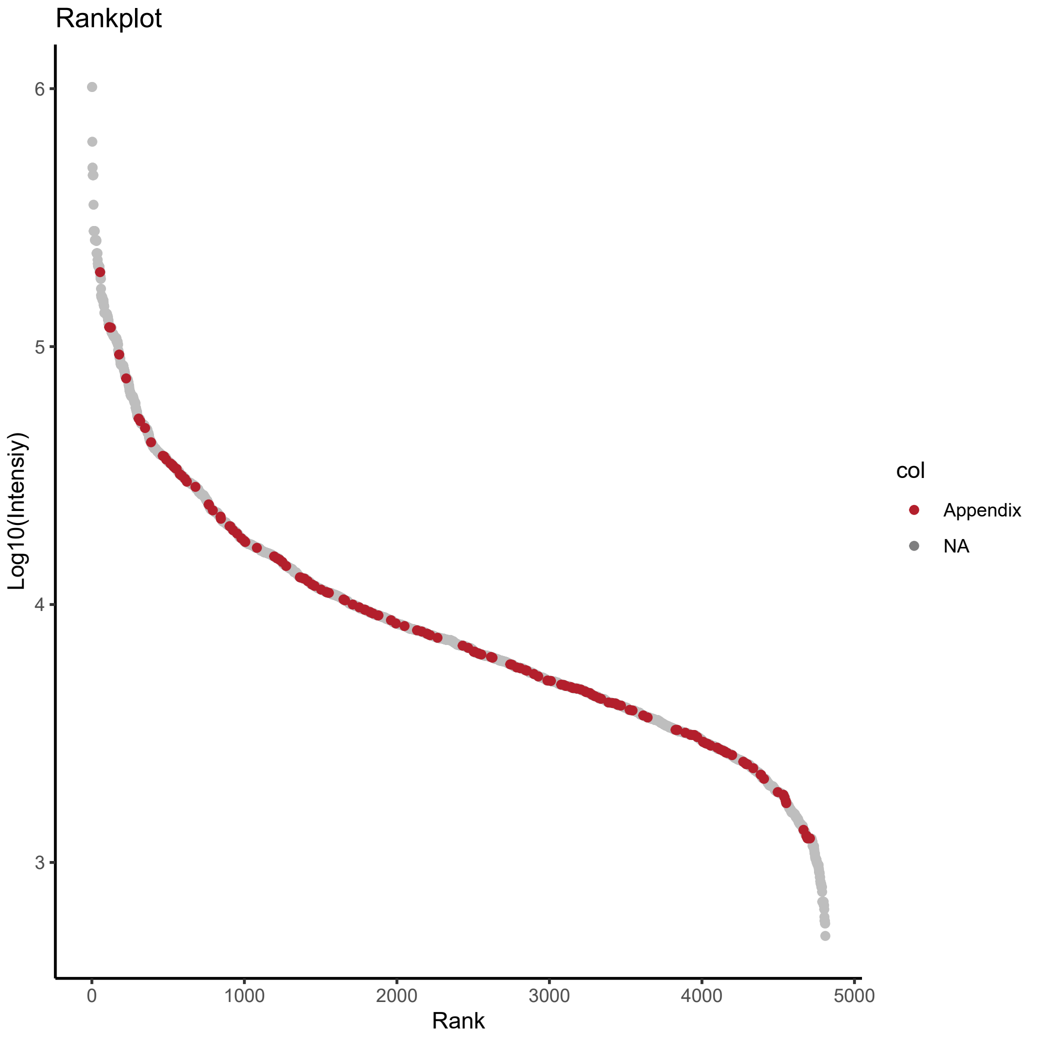
Rankplot
6
5
4
3
Log10(Intensiy)
0
1000
2000
3000
4000
5000
Rank
col
Appendix
NA

X)

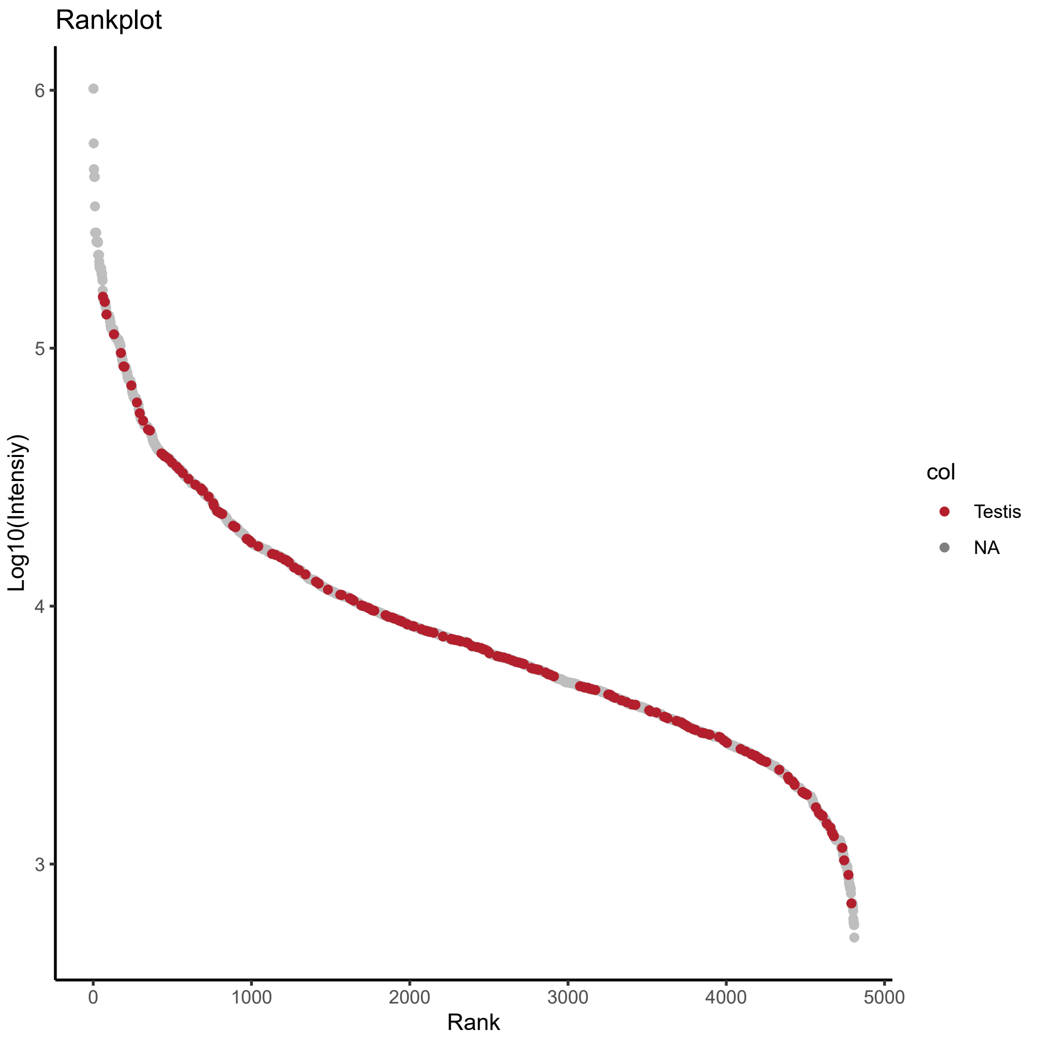
Rankplot
6
5
4
3
Log10(Intensiy)
0
1000
2000
3000
4000
5000
Rank
col
Testis
NA

Y)

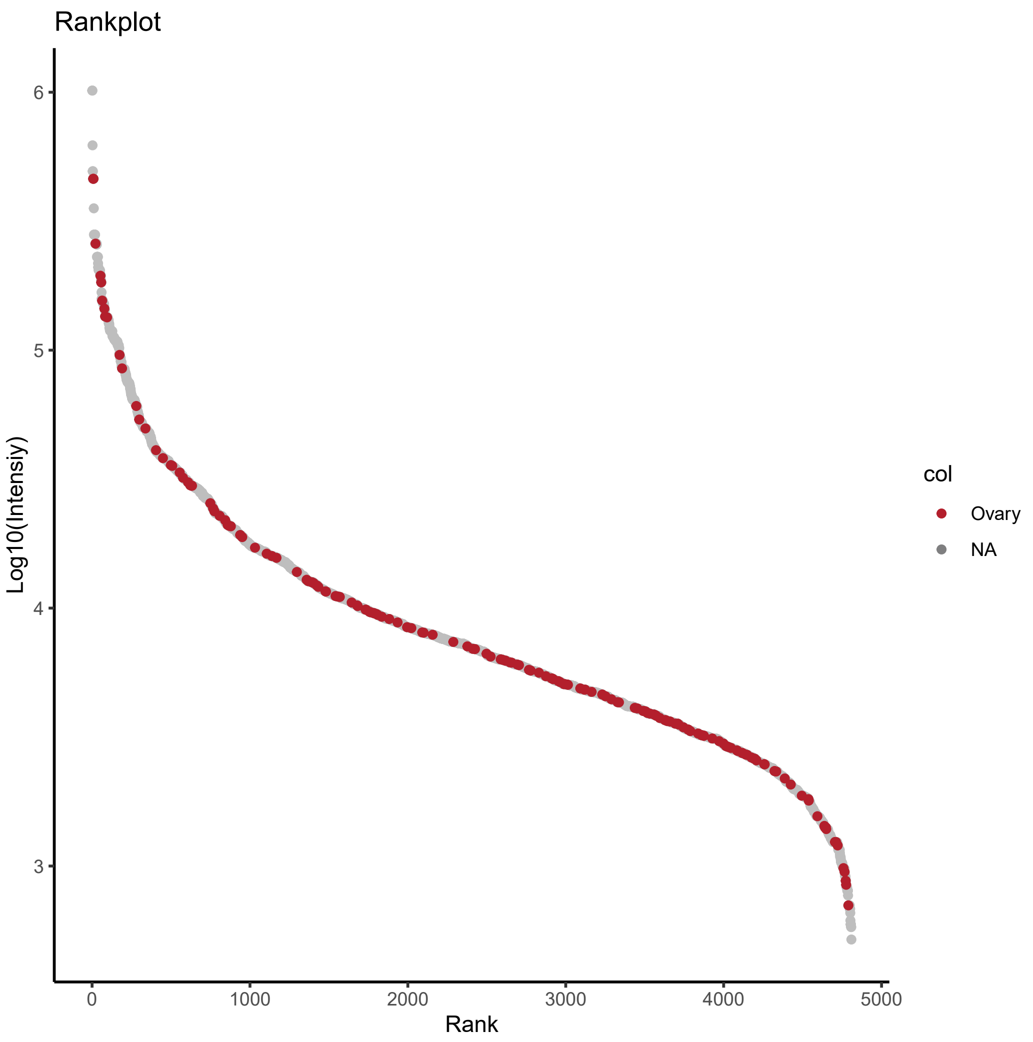
Rankplot
Log10(Intensiy)
6
5
4
3
0
1000
2000
3000
4000
5000
Rank
col
Ovary
NA

Z)

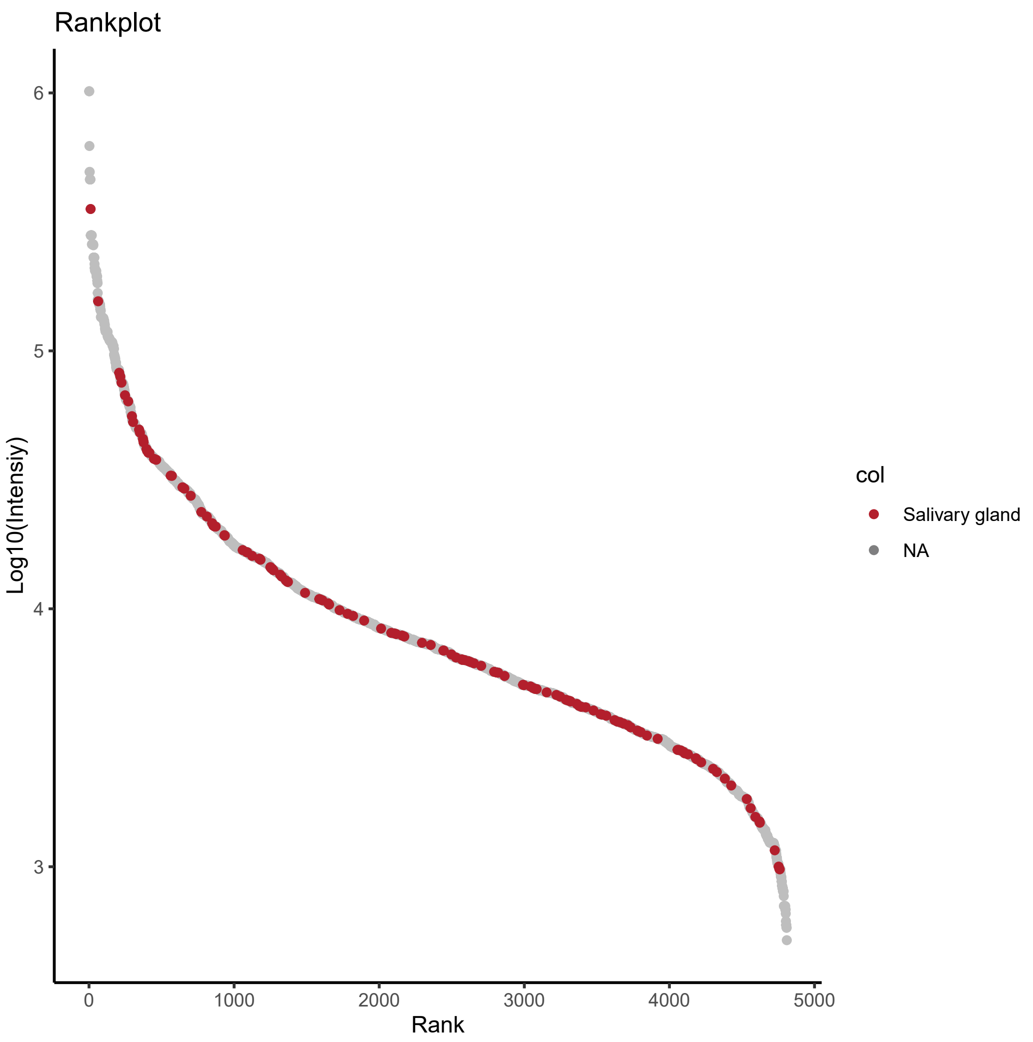
Rankplot
Log10(Intensiy)
6
5
4
3
0
1000
2000
3000
4000
5000
Rank
col
Salivary gland
NA

**Supplementary Figure 2: (A)-(Z)** Intensity rank plot of peptides derived from group enriched, tissue enhanced and tissue enriched genes colored by their respective organ assignment (dataset as in (Figure 4A)).
